# Non-synaptic exocytosis along the axon shaft and its regulation by the submembrane periodic skeleton

**DOI:** 10.1101/2025.09.17.676728

**Authors:** Theresa Wiesner, Christopher Parperis, Fanny Boroni-Rueda, Nicolas Jullien, Afonso Mendes, Léa Marie, Louisa Mezache, Marie-Jeanne Papandréou, Ricardo Henriques, Christophe Leterrier

**Affiliations:** Aix Marseille Université, CNRS, INP UMR7051, NeuroCyto, Marseille, France; Instituto de Tecnologia Química e Biológica António Xavier, Universidade Nova de Lisboa, Oeiras, Portugal; Gulbenkian Institute for Molecular Medicine, Oeiras, Portugal; UCL Laboratory for Molecular Cell Biology, University College London, London, UK

## Abstract

Neuronal communication relies on signaling molecules transferred via exo- and endocytosis throughout the brain. Historically, studies have focused on vesicle exo- and endocytosis and their release machinery at synapses, and much less is known about non-synaptic exocytosis. If and how vesicles can access the plasma membrane along the axonal shaft, overcoming the insulating layer of the membrane-associated periodic scaffold, remains unclear. Here, we used fast live-cell imaging of mature cultured hippocampal neurons expressing the vamp2-pHluorin reporter to map spontaneous exocytosis along axons. We detected non-synaptic exocytic events along the axon shaft that concentrated at the axon initial segment. Perturbation of the membrane-associated actin-spectrin skeleton revealed its role in regulating shaft exocytosis, similarly to its recently demonstrated role in shaping axonal endocytosis. To visualize the nanoscale arrangement of exocytic locations, we developed a novel correlative live-cell/two - color, single-molecule localization microscopy (SMLM) approach. We observed that regions of exocytosis are devoid of the submembrane spectrin mesh, with these spectrin-free areas being spatially separated from the spectrin clearings that contain clathrin-coated pits. Overall, our work reveals a new process of spontaneous exocytosis along the axon shaft, and how the axonal submembrane skeleton shapes a heterogeneous landscape that uniquely segregates vesicular trafficking events.

## Introduction

Neurons communicate by transmitting signals to target cells via the release of vesicles containing various molecules such as neurotransmitters, signaling, and trophic factors. A population of transport vesicles travels along the axon before reaching sites of communication (Guedes-Dias & Holzbaur, 2019), where vesicular delivery to and retrieval from the plasma membrane occurs by exo- and endocytosis (Mochida, 2022). For exocytosis, the majority of studies have focused on the release of neurotransmitter-filled vesicles at synapses (Ariel & Ryan, 2010; Jackson et al., 2022; Kavalali & Jorgensen, 2014; Leitz & Kavalali, 2014; Maschi & Klyachko, 2017; Südhof, 2004), but evidence exists of non-synaptic exocytosis outside of synaptic compartments along the axonal shaft of mature neurons in culture (de Wit et al., 2009; Ratnayaka et al., 2011a; van de Bospoort et al., 2012) as well as in the developing zebrafish spinal cord (Almeida et al., 2021). The relative scarcity of non-synaptic exocytosis observations likely reflects its lower occurrence, suggesting that specific mechanisms ensure preferential exocytosis in presynapses compared to the axon shaft.

Super-resolution microscopy has revealed a dense cytoskeletal assembly lining the plasma membrane of axons, composed of circumferential actin rings spaced every ~ 185 nm by spectrin tetramers (Leterrier et al., 2015; Xu et al., 2013). This scaffold is called the membrane-associated periodic skeleton (MPS, Zhong et al., 2014) and continuously runs along the axon shaft, stopping precisely at presynaptic boutons (He et al., 2016; Sidenstein et al., 2016). Furthermore, our laboratory recently revealed that along the proximal axon, the MPS restricts endocytosis within ~300 nm circular holes in the spectrin mesh – termed “spectrin clearings” (Wernert et al., 2024). The interruption of the MPS at presynaptic boutons, together with its restrictive role on endocytosis, suggest that the MPS could also regulate exocytosis along the axon shaft, forming an insulating layer that prevents transported vesicles from spuriously accessing the plasma membrane outside of synaptic compartments.

Supporting this hypothesis are studies in non-neuronal cell types proposing that cortical actin acts as a mesh under the plasma membrane, restricting vesicle exocytosis (for reviews see Miklavc & Frick, 2020; Porat-Shliom et al., 2013). Moreover, early work from the Aunis lab in chromaffin cells (Perrin et al., 1987) showed that spectrins inhibits calcium-induced exocytosis, suggesting an insulating layer function for the membrane-associated spectrin mesh (Aunis & Bader, 1988). While these works explore the role of the actin/spectrin scaffold in regulating membrane access, most studies in neurons have focused on the role of actin at presynapses, delineating its roles in organizing the synaptic vesicle pool and scaffolds (Glebov et al., 2017; Ogunmowo et al., 2023; Sankaranarayanan et al., 2003; Tumminia et al., 2024). A more specific study on non-synaptic exocytosis along axons showed actin recruitment following vesicle fusion at the diffraction-limited microscopy level (Ratnayaka et al., 2011a), but did not resolve further the environment of non-synaptic exocytic sites. If and how the stable MPS regulates and organizes exocytic events along the axon shaft remains unknown.

To address this, we first mapped exocytosis events along the axon and at presynapses using fast live-cell imaging of cultured hippocampal neurons expressing the exocytosis reporter vamp2-pHluorin. We observed a sparse occurrence of spontaneous exocytosis along the distal axonal shaft compared to nearby presynapses. We then assessed the amount of non-synaptic exocytosis occurring at distinct axonal compartments (axon initial segment AIS, proximal and distal axon) compared to the cell body and dendrites, and found that a larger fraction of spontaneous non-synaptic exocytosis occurs at the AIS. Acute perturbation of actin to disassemble the MPS resulted in an increase in shaft exocytosis, consistent with an insulating role. To understand the nanoscale context of shaft exocytosis, we developed a correlative strategy combining live-cell imaging of exocytosis followed by fixed-cell single molecule localization microscopy (SMLM). This revealed that exocytosis occurs within holes in the spectrin mesh, as previously observed for clathrin-mediated endocytosis taking place in spectrin clearings (Wernert et al., 2024). We further observed that exocytosis sites are mostly distinct from the spectrin clearings that contain clathrin-coated pits. Overall, our work highlights the underappreciated process of spontaneous axon shaft exocytosis and its regulation as a new physiological role of the MPS.

## Results

### Axonal shaft shows sparser exocytosis compared to presynapses

In mature neurons, the MPS is fully formed within the axon initial segment (AIS) and the proximal axon, with interruptions occurring at the presynapse (Figure 1A, Leterrier, 2021) and the distal axon (Boyer et al., 2024; Gazal et al., 2025). We first asked whether non-synaptic exocytosis can be observed along the axonal shaft (Ratnayaka et al., 2011b; van de Bospoort et al., 2012). We monitored spontaneous exocytosis events by performing HiLo microscopy in the axonal shaft of mature dissociated hippocampal neurons cultured for 12-16 days in vitro (DIV) transfected with a pH-sensitive fluorophore (pHluorin) tagged to the vesicle-associated membrane protein 2 (vamp2-pHluorin, Miesenböck et al., 1998; Sankaranarayanan et al., 2000). The protein vamp2 was chosen among the vesicular soluble N-ethylmaleimide-sensitive factor attachment protein receptors (SNAREs) as it is a key soluble SNARE protein involved in all forms of neurotransmission (Schoch et al., 2001; Wilhelm et al., 2014). Axonal regions were first illuminated at high laser power to photobleach pre-existing vamp2-pHluorin at the plasma membrane, maximizing the signal-to-noise ratio for the time-lapse fluorescent detection of exocytosis events (Figure 1B, Yudowski et al., 2007). Sites of constitutive exocytosis were identified by detecting pHluorin flashes using the recently developed pHusion software (O’Shaughnessy et al., 2024). We co-transfected the neurons with the presynaptic marker synaptophysin-mCherry (Navone et al., 1986; Wiedenmann & Franke, 1985) to classify the identified exocytotic events as synaptic (at presynapses) or non-synaptic (along the axonal shaft, Figure 1C-E, Movie S1).

**Figure 1:**
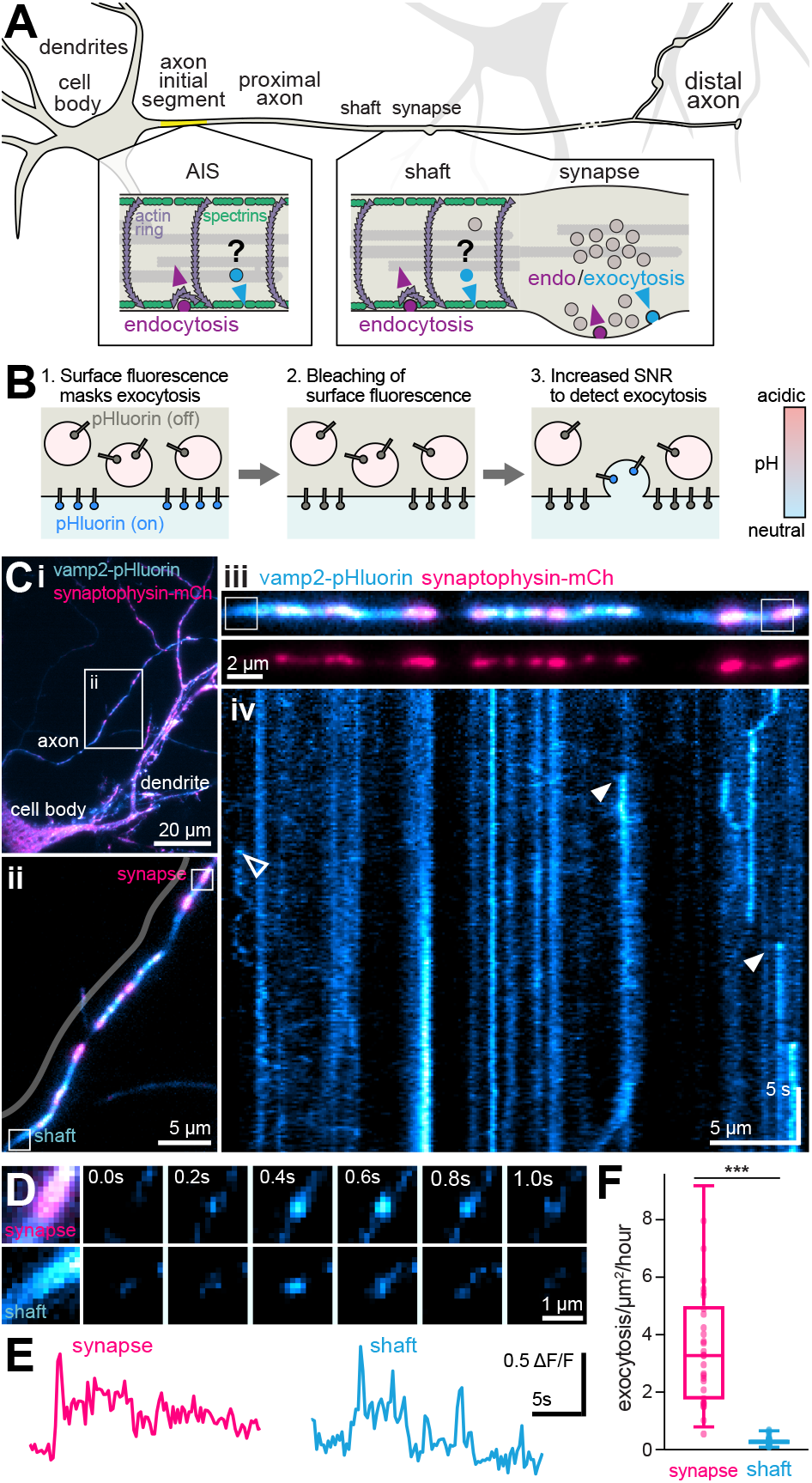
Spontaneous synaptic and non-synaptic exocytosis along the axon. **A**. Schematic neuron with its different neuronal compartments. The axonal compartment and underlying MPS, composed of actin rings interspaced by spectrin tetramers, are highlighted. While the plasma membrane is accessible to exo-endocytosis at the presynapse that is devoid of the MPS, vesicles within the axonal shaft might have restricted access to the plasma membrane due to the insulating role of the MPS. **B**. Schematic representation for the imaging of vesicular release using vamp2-pHluorin. **C**. i) Representative neuron co-expressing vamp2-pHluorin (cyan-hot) and synaptophysin-mCherry (pink). ii) Zoom of the axonal region shown in i) (white rectangle). iii) corresponding axon (parallel to the delineated line in ii) and iv) corresponding kymograph of vamp2-pHluorin dynamics (cyan-hot) showing exocytosis events. Full arrowheads indicate sites of exocytosis occurring at presynapses, empty arrowhead at axonal shaft. **D**. Zooms corresponding to the white rectangles shown in C ii). Shown are exocytosis events at a presynapse (positive for synaptophysin-mCherry, pink, top) and at the axonal shaft (bottom) with corresponding montage of vamp2-pHluorin time-lapses (cyan-hot). **E**. Fluorescence traces of vamp2-pHluorin during the exocytosis events occurring at the presynapse (left) or axonal shaft (right) pictured in E. **F**. Exocytosis rate (number of events per µm^2^ per hour) at presynapses or along the axonal shaft (n = 30-31 fields of view, 5 independent neuronal culture; Mann-Whitney test, for exact values refer to ***Supplementary Data***).

Compared to presynapses where spontaneous exocytosis occurred robustly (3.19 events/µm^2^/hour, all values in text are median unless specified otherwise), we detected ~20 times fewer exocytosis events along the axonal shaft (0.20 events/ µm^2^/hour; Figure 1F, refer to ***Supplementary Data*** for all exact values from graphs). These results indicate that although exocytosis occurs preferentially at presynapses, non-synaptic exocytosis can occur along the MPS-lined axonal shaft as previously reported (Ratnayaka et al., 2011b; van de Bospoort et al., 2012).

### The axon initial segment is a hotspot for non-synaptic exocytosis

Axons can be functionally and molecularly divided into the axon initial segment (AIS), proximal (region immediately after the AIS), and distal axon subcompartments (Leterrier et al., 2017). As we observed spontaneous non-synaptic exocytosis along the distal axonal shaft (hundreds of microns away from the cell body), we expanded our observations to the AIS and proximal axon, and compared it to non-synaptic exocytosis occurring at the cell body and the dendrites (devoid of the MPS) as well as distal axons visible in the same field of view (Figure 2A-B, Movie S2). Upon quantification, we found no significant difference between the exocytosis rate in dendrites (0.39 events/µm^2^/hour) and the cell body (0.26 events/µm^2^/ hour, Figure 2C). Interestingly, we observed robust exocytosis at the AIS (labeled for an extracellular epitope of the AIS-specific adhesion molecule neurofascin, Figure 2A-B, Movie S2), and measured a significantly higher rate of exocytosis (1.08 events/µm^2^/hour) in the AIS compared to the cell body or dendrites (Figure 2C). The exocytosis rate at the AIS was also significantly higher than in the proximal and distal axon (0.41 and 0.12 events/µm^2^/hour, respectively; Figure 2C). To test if the gradual attenuation of non-synaptic exocytosis could result from a difference of available t-SNARE proteins at the plasma membrane along the different axonal segments, we used Structured Illumination Microscopy (SIM) to resolve the distribution of syntaxin 1a along axons. We found no visible difference in the density of syntaxin clusters, confirming its presence along the whole axon (Figure S1, Maidorn et al., 2018). These experiments show that non-synaptic exocytosis along the proximal and distal axonal shaft is similar in density to exocytosis along the cell body and dendrites, with the AIS being a hotspot of exocytosis.

**Figure 2:**
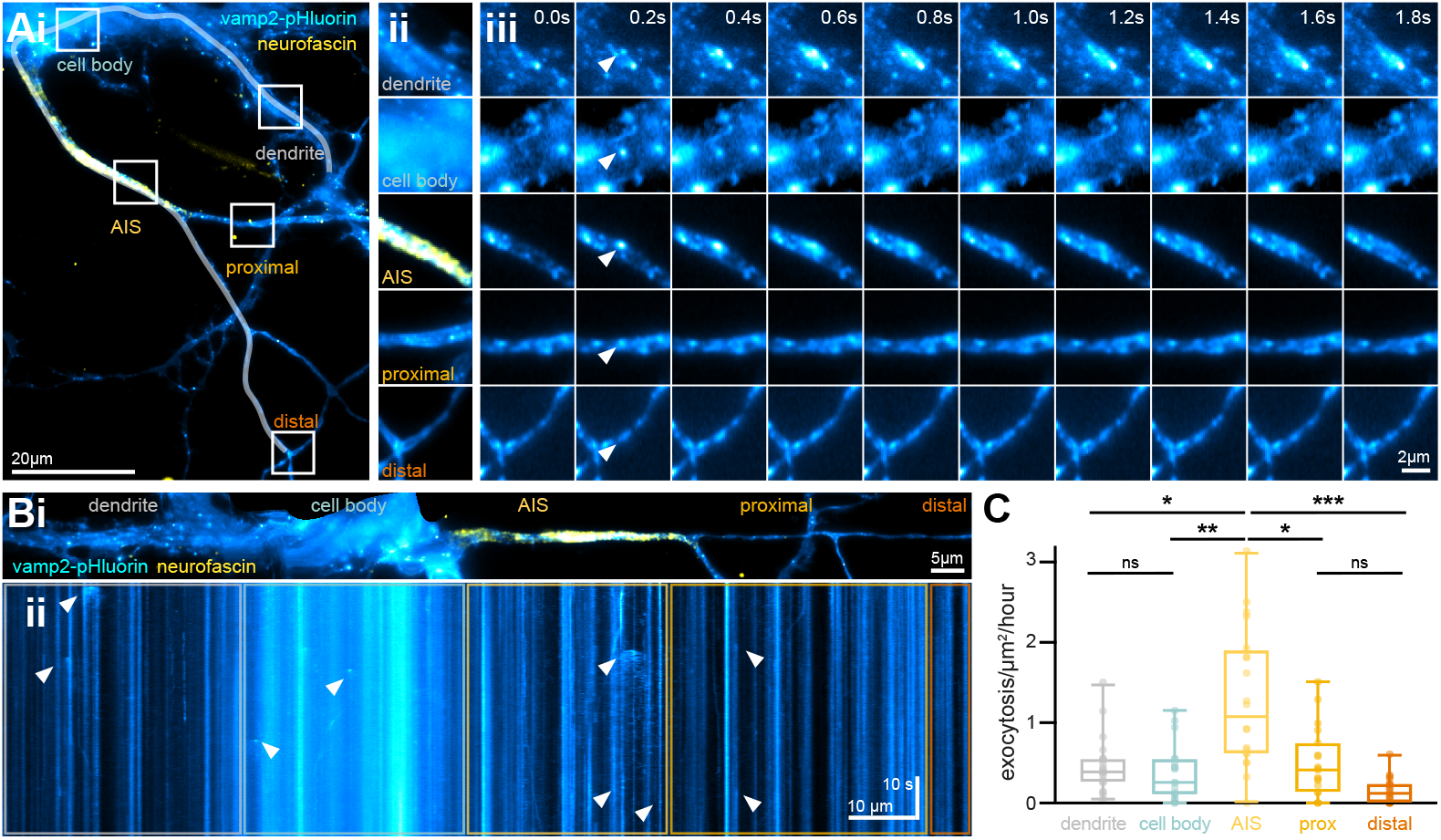
Mapping of exocytosis in different neuronal compartments. **A**.i) Representative neuron expressing vamp2-pHluorin (cyan-hot) with the AIS labeled for neurofascin (yellow). White rectangles indicate different neuronal compartments where exocytosis events are observed: dendrites, cell body, AIS, proximal, and distal axon. ii) Zooms on the area highlighted by white rectangles in i). iii) corresponding montage of exocytosis events (cyan-hot). Arrowheads (white) indicate sites of exocytic events. **B**. i) Straightened area highlighted by the white line in i) showing a dendrite, the cell body, axon initial segment, proximal and distal axon. (ii) corresponding kymograph of vamp2-pHluorin dynamics (cyan-hot). Arrowheads (white) indicate sites of exocytosis events. **C**. Rate of exocytosis in: dendrites (20 fields of view), cell body (19 fields of view), AIS (20 fields of view), proximal (19 fields of view), and distal axon (20 fields of view, 3 independent cultures; ANOVA with Kruskal-Wallis test).

### Perturbation of the MPS increases non-synaptic exocytosis along the axon

We observed sparse non-synaptic exocytosis events along the axonal shaft, suggesting that the MPS restricts exocytosis by limiting access to the plasma membrane. To test this hypothesis, we acutely perturbed the MPS by applying the actin-severing drug swinholide A (swinA, Spector et al., 1999). In line with previous works (Mendes et al., 2025; Vassilopoulos et al., 2019; Wernert et al., 2024), swinA treatment resulted in a disappearance of filamentous actin staining and a disordering of the spectrin periodicity observed by SIM both along the AIS, proximal, and distal axon (Figure 3A). We then examined the rate of exocytosis events after swinA treatment and observed a significant increase in the exocytosis rate at the AIS (vehicle: 0.64 events/µm^2^/hour; swinA: 1.32 events/µm^2^/hour, Figure 3B). The increase in accessible bare membrane proportion at the AIS caused by swinA treatment (Wernert et al., 2024) thus leads to an increase in exocytic events, suggesting an insulating role for the MPS.

**Figure 3:**
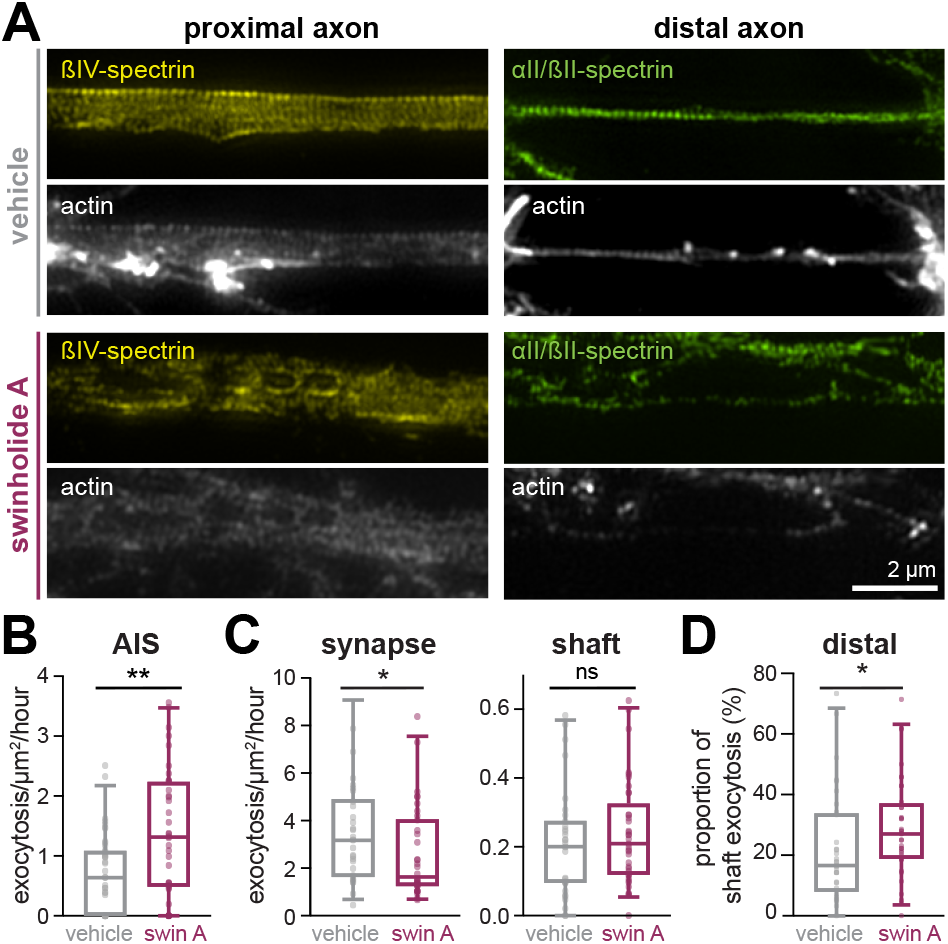
Modulation of exocytosis by acute perturbation of the MPS. **A**. Representative maximum projection images acquired using structured illumination microscopy (SIM) of βIV-spectrin (yellow), αII/βII-spectrin (green) and filamentous actin (grey) treated with either vehicle (grey) or swinA (100 nM of 3 h, purple) in the proximal (left) or distal axon (right). **B-C**. Rate of exocytosis events in vehicle or swinA-treated cultures at the AIS (B, fields of view = 43,37; 4 independent cultures); distal axonal shaft (C, left, fields of view = 31, 37; 5 independent cultures); and presynapse (C, right, fields of view = 30, 34; 5 independent cultures). D) Proportion of axonal shaft exocytosis events in vehicle or swinA-treated cultures (fields of view = 33, 36; 5 independent cultures; Mann-Whitney test for all graphs).

In distal axons as a whole, we observed no significant change in the spontaneous exocytosis rate after swinA treatment (vehicle: 0.65 events/µm^2^/hour, swinA: 0.58 events/µm^2^/ hour, Figure S2). However, when we classified the observed exocytic events along the distal axon into synaptic or non-synaptic, we observed a significant decrease in the exocytosis rate at presynapses (vehicle: 3.19 events/µm^2^/hour, swinA: 1.65 events/µm^2^/hour, Figure 3C), in line with a facilitating role of presynaptic actin for vesicular release (Bingham et al., 2023). By contrast, the exocytic rate along the axonal shaft was stable (vehicle: 0.20 events/µm^2^/hour, swinA: 0.21 events/µm^2^/ hour, Figure 3C). This led the proportion of non-synaptic exocytosis over total exocytosis (non-synaptic and synaptic over the whole area) to increase upon swinA treatment (vehicle: 16.7%, swinA: 27.1%, Figure 3D). This indicates that in the distal axon, actin perturbation impedes synaptic exocytosis as previously shown for stimulated release (Ganguly et al., 2015) and favors non-synaptic exocytosis along the shaft.

### Exocytosis occurs at spectrin-less membrane areas

We recently demonstrated that in the AIS and proximal axon, endocytosis occurs in ~300-nm circular areas that are devoid of spectrin and termed these regions spectrin clearings. These spectrin clearings allow for the formation of stable clathrin-coated pits (CCPs) on the bare plasma membrane at their center as the first step of a regulated endocytosis process (Wernert et al., 2024). We thus wondered if the axonal shaft exocytosis would also need to occur in “holes” of the spectrin mesh, and if these were the same clearings used for CCP formation. To test this, we developed a correlative live-cell/SMLM approach. We first mapped sites of vamp2-pHluorin exocytosis by HiLo microscopy in living neurons as above, then fixed and labeled them for both αII- and βII-spectrin, and performed SMLM to image the nanoscale arrangement of spectrins along the imaged axon (Figure 4A & 4F). We aligned the live-cell imaging data to the reconstructed SMLM images to reveal the nanoscale environment of exocytic sites (Figure 4B & 4G). To quantify the periodicity of αII- and βII-spectrins on SMLM images, we used the recently developed Structural Repetition Detector approach (SReD; Mendes et al., 2025).

**Figure 4:**
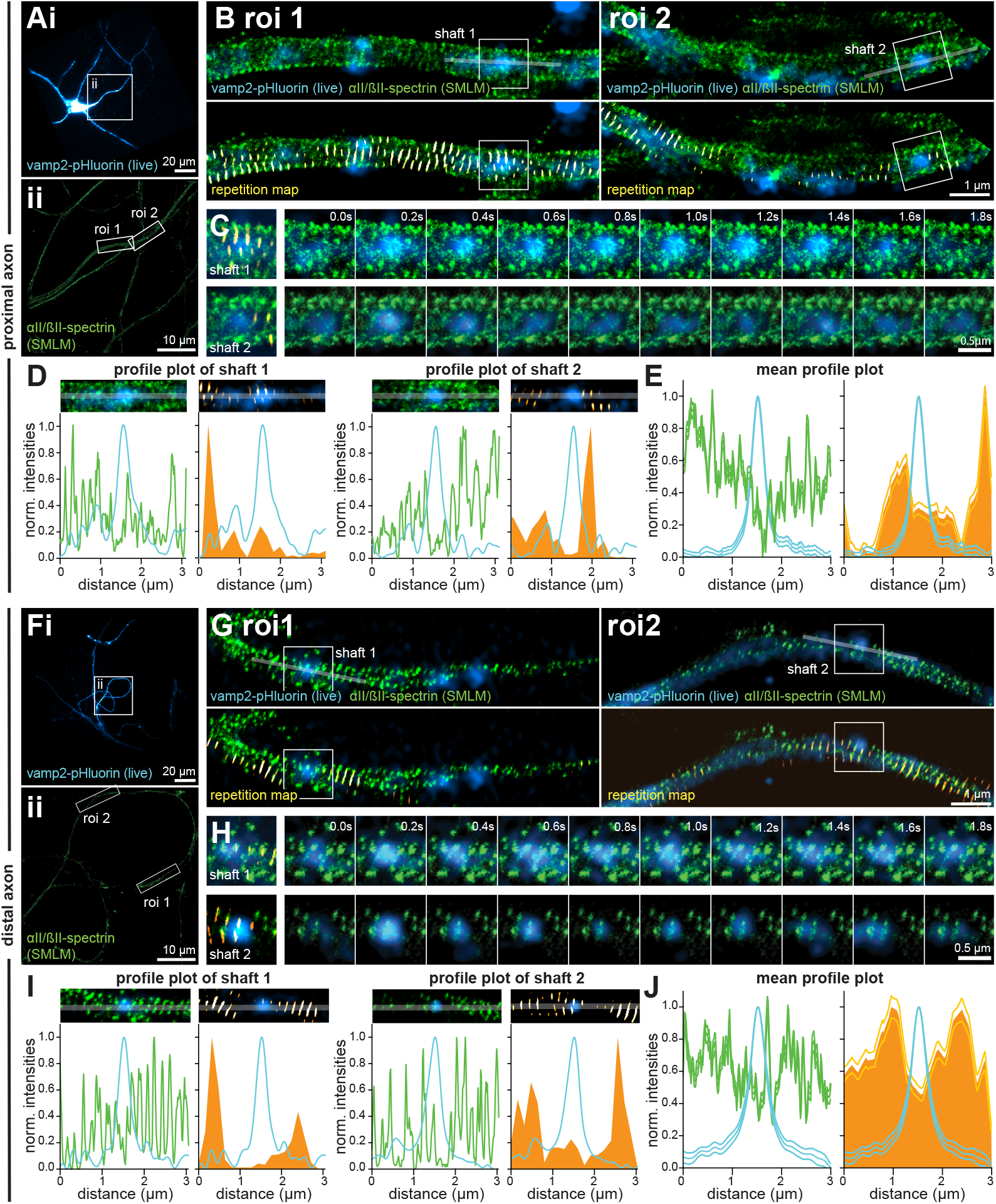
Axonal shaft exocytosis occurs at sites devoid of spectrin mesh. **A and F**. (Top) Representative neurons expressing vamp2-pHluorin (cyan), which have been affine transformed to align with (bottom) the corresponding SMLM images of αII/βII-spectrin (green). White rectangles in A and F (top) delineate the regions acquired with the SMLM microscope. **B and G**. Regions of interest shown in A and F, bottom. Top, Overlays of vamp2-pHluorin (cyan) and αII/βII-spectrin (green). Bottom, additional overlay of the structural repetition map generated by SReD (hot-orange). **C and H**. Regions of exocytosis events shown in B and G. Left, first frame of exocytosis (hot-cyan), corresponding αII/βII-spectrin SMLM images (green), and repetition map (hot-orange). Right, Montages of the exocytic event (hot-cyan) and corresponding αII/βII-spectrin SMLM image (green). Exocytic event starts at 0.2 seconds. **D and I**. Representative images (top) and profile plots (bottom) of axonal segments 1 and 2 shown in B and G. Shown are the profile plots of the normalized intensity of the exocytic event (cyan), αII/βII-spectrin (green, left) and the repetition score (orange, right). **E and J**. Average profile plots of exocytic events (cyan), αII/βII-spectrin labeling (green, left), and repetition score (orange, right). Average profiles were calculated from 34 exocytosis events of 5 neurons from 4 independent cultures (proximal axon, E) and 26 exocytosis events of 4 neurons, 2 independent cultures (distal axon, J). Surrounding lines are the associated SEM values.

Along the proximal and distal axonal shaft, we observed that most exocytic events occurred in sites either devoid of αII/βII-spectrin, or where the αII- and βII-spectrin periodicity was disrupted (Figure 4C & 4H, Movie S3 & S4). Indeed, intensity profiles and SReD detection showed exocytic events occurring in areas of low αII- and βII-spectrin intensity and low repetition score (Figure 4D & 4I). Quantification of the average αII/βII-spectrin intensity as well as the average repetition score confirmed that exocytic sites were sites of low spectrin intensity and periodicity compared to their surroundings (Figure 4E & 4J). These results indicate that exocytosis requires regions of spectrin-less plas-ma membrane along the axon shaft, in line with an insulating role of the MPS.

### Exocytic sites are distinct from CCP-containing spectrin clearings

Having shown that exocytosis occurs in holes of the spectrin mesh along the axonal membrane, we next wanted to know if these holes could be the spectrin clearings where CCP formation and regulated endocytosis occur (Wernert et al., 2024), or if the two processes were spatially segregated in distinct spectrin holes. To address this question, we expanded our correlative live-cell/SMLM approach to multicolor SMLM by coupling live-cell with spectral demixing stochastic optical reconstruction microscopy (STORM, Figure 5A & 5G; Friedl et al., 2023). This allowed us to image the nanoscale arrangement of spectrins around exocytic sites together with spectrin clearings that contain CCPs (Figure 5B & 5H).

**Figure 5:**
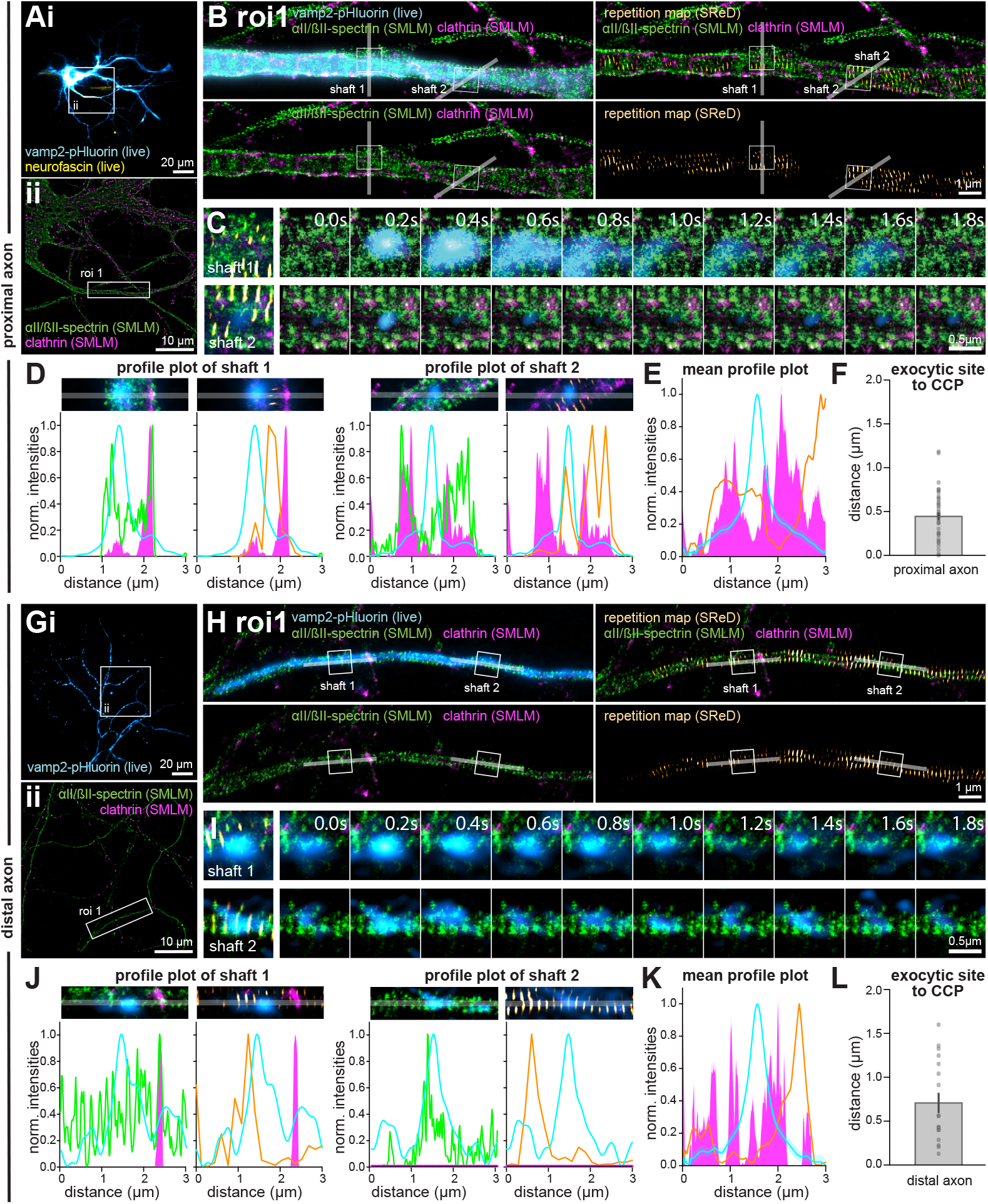
Axonal exocytosis and endocytosis occur in distinct spectrin-less areas. **A and G**. Top, representative neurons expressing vamp2-pHluorin (cyan in A), with their AIS labeled for neurofascin (yellow). Bottom, corresponding aligned SMLM images of αII/βII-spectrin (green) and clathrin (magenta). White rectangles in A and G (top) delineate the regions acquired with SMLM. **B and H**. Regions of interest shown in A and G, bottom. Top left, overlays of vamp2-pHluorin (cyan), αII/βII-spectrin (green), and clathrin (magenta). Bottom left, super-resolved images of αII/βII-spectrin (green) and clathrin (magenta). Top right, overlay of vamp2-pHluorin (cyan), αII/βII-spectrin (green), clathrin (magenta), and SReD repetition map (hot-orange). Bottom right, SReD repetition map alone (hot-orange). **C and I**. Regions of exocytosis events shown in B and H. Left, first frame of the exocytic event (hot-cyan), corresponding αII/βII-spectrin (green) and clathrin (magenta) SMLM images, and SReD repetition map (hot-orange). Right, montages of exocytic events (hot-cyan) with corresponding αII/II-spectrin (green) and clathrin (magenta) SMLM images. Exocytic event starts at 0.2 seconds. **D and J**. Representative images (top) and profile plots (bottom) of axonal segments 1 and 2 shown in B and H. Shown on the plot are the normalized intensity of the exocytic event (cyan), αII/βII-spectrin (green, left), clathrin (magenta, left), and the repetition score (orange, right). **E and K**. Average profile plots of exocytic events (cyan), clathrin (magenta), and SReD repetition score (orange). Surrounding lines are the associated SEM values. **F and L**. Distances between exocytic sites and nearest clathrin clusters. Average profiles and distances were calculated from 39 exocytosis events of 5 neurons, 4 independent cultures (AIS and proximal neuron, E and F) and 17 exocytosis events of 4 neurons from 2 independent cultures (distal axon, K and L).

As with single-color SMLM above, we observed that exocytic events occur in regions of low spectrin intensity and low periodicity (Figure 5C & 5I, Movies S5 and S6). In addition, we observed that CCPs were often found in spectrin clearings adjacent to the precise site of exocytosis, and not directly present in the same spectrin-less area (Figure 5C & 5I, Movies S5 and S6). Intensity profiles confirmed the spatial separation of the clathrin clusters from the exocytic site, even if both occur in regions of low spectrin intensity and repetition score (Figure 5D-E & 5J-K). We quantified the average distance between the center of exocytic sites and the nearby clathrin clusters in the proximal and distal axon to 460 nm and 720 nm, respectively (Figure 5F & L). The complexity of the correlative live-cell/ SMLM imaging makes it difficult to acquire a large number of neurons, but the larger distance between exocytic sites and surrounding CCPs was nevertheless significant when comparing the proximal to the distal axon (Figure S3). In line with this, in the proximal axon, 2.5% of exocytic sites showed no adjacent CCP within a 3 µm distance, whereas this proportion was 34.6% for the distal axon, indicating a larger separation between exocytic and endocytic sites in the latter. Finally, we confirmed the separation between exocytic and endocytic sites by examining our SIM data of t-SNARE syntaxin 1A along axons, showing syntaxin clusters in holes of the spectrin mesh, but spatially distinct from CCPs at the AIS and along the distal axon shaft (Figure S1).

Our correlative results show that exocytic sites are not occupied by a stable CCP within 5-15 minutes of exocytic event detection, considering the acquisition and fixation time. To gain a better insight into the relative dynamics of exocytosis and CCP formation and detect a possible succession of endocytic and exocytic event at the same site, we next performed HiLo imaging of living neurons co-expressing vamp2-pHluorin and clathrin-mCherry (Figure 6A). To better capture exocytic events, we used a combination of fast vamp2-pHluorin imaging (15 seconds of continuous imaging) interspersed with snapshots of clathrin-mCherry for a total duration of 5 minutes. We observed that exocytic sites generally did not overlap with CCPs (Figure 6A, Movie S7), with intensity profiles of the exocytic event (vamp2-pHuorin from the first frame of the event) and the surrounding CCPs (from the closest acquisition time point) showed a spatially distinct distribution (Figure 6B). This was confirmed by calculating the average profile plots of the exocytosis events and clathrin clusters (Figure 6C). The average distance between the exocytic site and the surrounding clathrin clusters was approximately 1 μm (Figure 6D). We then classified sites of exocytosis into 3 categories: i) sites exclusively used by exocytosis (no overlap with a CCP over the 5-minute acquisition); ii) sites of sequential use of exocytosis and clathrin clusters (detection of both exocytic event and CCP during the 5-minute acquisition); and iii) sites of simultaneous occurrence of exocytosis and CCP (on the same frame). We found that the majority of exocytic sites were used exclusively for exocytosis (mean 69.2%), while around one quarter saw the non-simultaneous appearance of CCP at the same location (23.3%). Simultaneous exocytosis and CCP presence was observed in only a minority of sites (7.5%, Figure 6E & 6F). These results confirm that spontaneous exocytosis and endocytosis occur at distinct spectrin-less sites along the axon.

**Figure 6:**
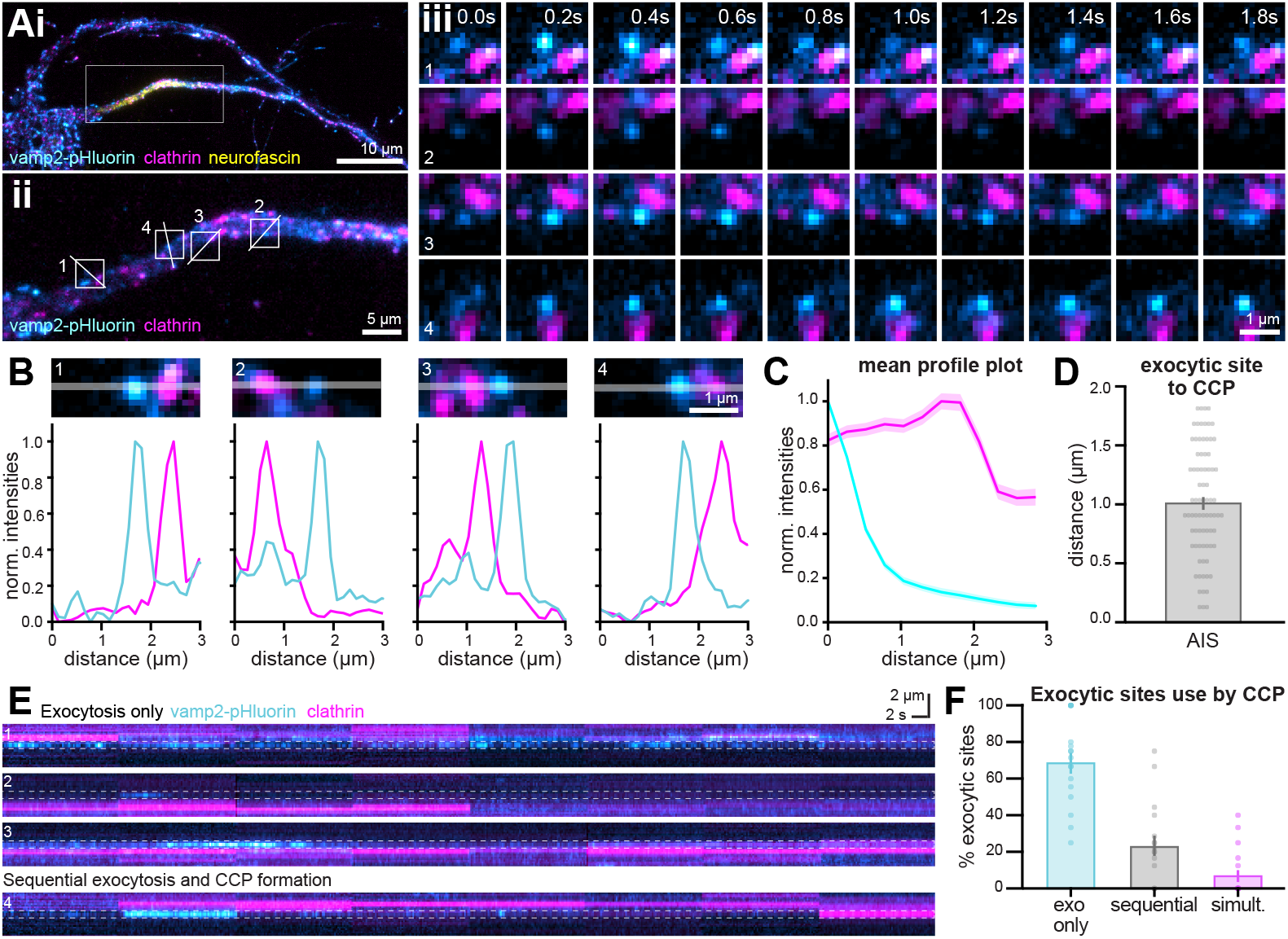
Exocytosis and endocytosis occur at distinct sites spatially and temporally. **A**. i) Representative live neuron expressing vamp2-pHluorin (cyan-hot) and clathrin-mCherry (magenta). The AIS was labeled for neurofascin (yellow). White rectangle delineates regions shown in iii). ii) Zoom corresponding to the white rectangle of the top image. iii) Montages of regions of interest shown in ii). Exocytic events start at 0.2 seconds. **B**. Representative images (top) and profile plots (bottom) of regions of interest drawn in ii). Shown are the profile plots of the normalized intensity of the exocytosis event (cyan) and the nearest clathrin cluster (magenta). **C**. Average profile plots of exocytosis sites (cyan) and nearest clathrin cluster (magenta). Average profiles were calculated from 79 exocytosis events of 17 neurons from 4 independent cultures. Surrounding lines are the associated SEM values. **D**. Distance between exocytic sites and the nearest clathrin clusters (exocytosis events = 79, neurons = 17, 4 independent cultures). **E**. Kymographs of vamp2-pHluorin (cyan-hot) and clathrin-mCherry (magenta) showing exocytosis sites exclusively used by exocytosis (kymographs 1-3) and sites sequentially used by exocytosis (blue horizontal signal) or CCPs (magenta horizontal signal, kymograph 4). Dashed lines indicate the site of exocytosis throughout time. **F**. Proportion of exocytic sites categorized by CCP occupancy. Data obtained from 17 neurons from 4 independent cultures.

## Discussion

While the majority of studies on axonal exocytosis focus on synaptic release at presynapses (Ariel & Ryan, 2010; Jackson et al., 2022; Kavalali & Jorgensen, 2014; Leitz & Kavalali, 2014; Maschi & Klyachko, 2017; Südhof, 2004), few studies describe non-synaptic exocytosis along the axon shaft (Almeida et al., 2021; De-Miguel et al., 2021; Ratnayaka et al., 2011b; van de Bospoort et al., 2012). How non-synaptic vesicles access the plasma membrane at the axonal shaft, where they have to overcome the insulating barrier of the MPS, remains elusive.

We confirm that non-synaptic exocytosis can occur at the axonal shaft as was previously observed (de Wit et al., 2009; van de Bospoort et al., 2012). We observe a low occurrence of spontaneous non-synaptic vesicular release, in line with what had been observed for dense core vesicle release in mature hippocampal neurons (de Wit et al., 2009; van de Bospoort et al., 2012), although in these cases exocytosis had been electrically stimulated. Interestingly, we find axonal shaft exocytosis to occur prominently at the AIS. Previous work in developing neurons has shown that insertion of the transmembrane protein neurofascin and voltage-gated potassium channel in the AIS occurs by lateral diffusion into the AIS from exocytosis occurring at the cell body and nerve terminal (Ghosh et al., 2020). However, studies in more mature neurons show that voltage-gated sodium and potassium channels get inserted via exocytosis at the AIS (Akin et al., 2015; Yoshioka & Okamura, 2024). These observations are in line with our results, and may indicate a developmental shift in membrane protein insertion into the AIS from lateral diffusion to localised exocytosis.

Exocytosis along the axon shaft requires access to the plasma membrane, raising questions about its regulation by the MPS and its dense mesh of spectrins lining the plasma membrane. We show that non-synaptic exocytosis at the AIS is restricted by the MPS, as we observe an increase of the exocytosis rate after acute MPS perturbation. This is similar to the effect of MPS perturbation on the formation of CCPs, due to increased membrane access (Wernert et al., 2024). In the distal axon beyond the AIS, we observed a smaller effect, with a proportional increase in non-synaptic exocytosis events, but no significant increase in exocytosis rate. Moreover, we measured a lower spontaneous exocytosis rate in control conditions at the distal axon compared to the AIS and proximal axon. A decrease in vesicular trafficking is expected in more distal portions of the axon due to cumulative delivery along its length, and could contribute to this effect (Bressloff, 2021).

More specific mechanisms might also regulate vesicular release along the distal axon shaft, such as a distinct composition and distribution of the exocytic machinery. We aimed to detect the majority of exocytosis types in mature neurons by using the v-SNARE protein vamp2, which is highly expressed throughout the neuronal development (Schoch et al., 2001; Wilhelm et al., 2014) and involved in a wide range of exocytosis types: plasma membrane addition in neuronal morphogenesis (Urbina et al., 2018; Urbina & Gupton, 2020), vesicle release of synaptic proteins (Bakr et al., 2021), release of dense core vesicles (Shimojo et al., 2015), and exosome release (Vilcaes et al., 2021) across the different neuronal compartments (Bakr et al., 2021; Yan et al., 2022). It is nonetheless possible that a fraction of the overall exocytosis along the distal axon is supported by different v-SNAREs. The v-SNARE vamp7 is mainly characterized in immature neurons, where it participates in neurite outgrowth (Burgo et al., 2012; Wojnacki et al., 2020), and has been shown to be present on lysosomes and a small proportion of vesicles in the synaptic vesicle pool of mature neurons (Hua et al., 2011; Ramirez et al., 2012, Proux-Gillardeaux et al., 2007). Yet, the exocytosis rate of vesicles containing vamp7 is generally lower than that of other v-SNARE coated vesicles (Bakr et al., 2021; Urbina et al., 2018b). Conversely, the difference in shaft exocytosis might involve a distinct distribution of t-SNARE proteins. However, we and others found the t-SNARE protein syntaxin 1a to be distributed at the distal axonal compartment (Figure S3, Maidorn et al., 2018). Finally, this difference might reflect the relative concentration of exocytic proteins at presynapses along the distal axon to support synaptic release, resulting in less nearby shaft exocytosis compared to the AIS and proximal axon that are largely devoid of presynaptic boutons.

Our correlative live-cell/SMLM approach allowed us to zoom in on the organization of shaft exocytosis. We found these sites to be spectrin-less membrane areas that are distinct from the spectrin clearings occupied by CCPs (Wernert et al., 2024). Whether the spectrin-less exocytic sites correspond to the ~15% of spectrin clearings that are unoccupied by CCPs (Wernert et al., 2024) or to distinct nanoscale holes in the MPS remains an open question: the precise identification of spectrin clearings requires platinum replica electron microscopy of unroofed axons, but exocytic sites are not readily visible on PREM views, in contrast to sites of clathrin-mediated endocytosis. It remains also unclear whether these exocytic holes are stable like the CCP-containing clearings, or if they represent more transient openings in the MPS as recently reported (Heller et al., 2025). To answer this question a fast, dynamic, live-cell, multicolor, super-resolution approach would be needed to simultaneously observe exocytosis events and the spectrin mesh organization over a long period of time. Nonetheless, it is clear that exocytosis and CCP formation occur at distinct locations. While our correlative live-cell/SMLM approach results in a ~5-20 minutes delay between the end of the live-cell imaging and the fixed cell labeling, it is fast enough to correlate exocytosis sites with the underlying stable spectrin mesh (Leterrier et al., 2015; Zhong et al., 2014) as well as with the slow-remodeling clathrin-coated pits (Wernert et al., 2024). Furthermore, live-cell imaging of exocytosis together with CCPs confirm that there is little spatiotemporal overlap between the two processes, with only a quarter of the exocytotic sites being used at different times for CCP formation.

What content is released by non-synaptic exocytosis we observe remains speculative at this point. In neurons, an alternative communication pathway to synaptic transmission is volume transmission, which allows the distribution of a variety of signaling molecules: neurotransmitters (dopamine, serotonin, acetylcholine…), growth factors (such as BDNF) and peptides (NPY released by dense core vesicles) (De-Miguel et al., 2021). Furthermore, during neuronal development, axons have also been shown to release synaptic organizers via lysosomes and autophagosomes (Cuhadar et al., 2024; Ibata & Yuzaki, 2021; Pascual-Caro & Juan-Sanz, 2024) to initiate and build the synaptic contact sites between neurons. Extrasynaptic communication between axon and oligodendrocyte has also been reported (Almeida et al., 2021) as well as the release of extracellular vesicles (Vilcaes et al., 2021) for intercellular communication. The molecular identity of the exocytosed cargoes remains to be investigated, notably at the AIS, and will shed further light on how the MPS shapes intercellular communication. So far, our work shows that the MPS shapes the pattern of exocytosis along the axon, defining sites of non-synaptic vesicle release and organizing a heterogeneous nanoscale landscape that segregates vesicular trafficking events. Disorganization of the MPS may create abnormal patterns of exocytosis along the axon that are likely to have significant consequences on the building and maintenance of the axonal arborization, and subsequent intercellular communication (Huang et al., 2017; Lorenzo et al., 2023).

## Acknowledgments

We thank the Gupton laboratory for sharing the plasmid vamp2-pHluorin and the Roy laboratory for sharing the clathrin-mCherry. We would like to acknowledge funding by the Agence Nationale pour la Recherche (ANR-20-CE13-0024 to C.L., ANR-21-CE42-0015 to C.L., ANR-20-CE16-0021 to C.L.), Fondation pour la Recherche Médicale (EQ202103012966 to C.L.), Fédération pour la Recherche sur le Cerveau (AOE 16 “Espoir en tête” 2021 to C.L.), and to thank the Neuro-Cellular Imaging Service and Nikon Center for Neuro-NanoImaging at INP, funded by CPER-FEDER (PlateForme NeuroTimone PA0014842). The project leading to this publication has received funding from Excellence Initiative of Aix-Marseille University - A^*^MIDEX, a French “Investissements d’Avenir” programme (AMX-19-IET-002 to T.W.). A.M.M and R.H. acknowledge support from the European Research Council (ERC) under the European Union’s Horizon 2020 research and innovation programme (grant agreement No. 101001332) and funding from an European Innovation Council (EIC) Pathfinder Open grant. However, the views and opinions expressed are those of the authors only and do not necessarily reflect those of the European Union. Neither the European Union nor the granting authority can be held responsible for them. This work was also supported by a European Molecular Biology Organization (EMBO) installation grant (EMBO-2020-IG-4734 to R.H.), and a Chan Zuckerberg Initiative Essential Open-Source Software for Science (EOSS6-0000000260 to R.H.).

## Author Contributions

TW performed the live-cell imaging. NJ devised the constructs used for live-cell imaging. TW, CLP, FB and LM prepared the immunostainings samples. FB prepared neuronal cell culture. CLP and TW performed the SMLM imaging. TW analyzed the exocytosis videos. CLP analyzed the SMLM images. AM, RH implemented SReD. AM and TW analyzed the SMLM data with SReD. TW performed the correlative data analysis with help from LM. TW wrote the original manuscript with guidance of CL. TW, CLP, FB, LM, AM, MJP, RH and CL edited the manuscript. TW, CL, MJP planned the experimental design. CL supervised the project.

## Methods

### Sample preparation

#### Neuronal culture

Primary neuronal cell culture was obtained in a similar procedure as previously described (Bingham et al., 2023). Briefly, hippocampi were extracted from E18 rat embryos from pregnant female Wistar rats (Janvier labs), dissected, and homogenized by trypsin treatment followed by mechanical trituration and seeded on 18-mm diameter round, #1.5H coverslips at a density of either 6,000 cells/cm^2^ (for correlative live-cell/ SMLM experiments) or 12,000 cells/cm^2^ (for the remaining experiments) for 3 hours in serum-containing plating medium (MEM with 10% fetal bovine serum, 0.6% added glucose, 0.08 mg/mL sodium pyruvate, 100 UI/mL penicillin-streptomycin). In accordance with the Banker method (Kaech & Banker, 2006), the coverslips (cells facing down) were then transferred to and cultured in petri dishes containing confluent astrocyte cultures conditioned in NB+ medium (Neurobasal medium supplemented with 2 % B-27, 100 UI/mL penicillin/ streptomycin and 2.5 µg/mL amphotericin). All procedures were in agreement with the guidelines established by the European Animal Care and Use Committee (86/609/ CEE) and was approved by the local French ethics committee (agreement G13O555).

#### Neuronal transfection

12-14 DIV neurons were transfected using Lipofectamine 2000 (Invitrogen, Ref 11668-027) with 0.25 μg of vamp2-pHluorin plasmid (gift from the Gupton laboratory, Urbina, et al., 2018) and either 0.25 μg synaptophysin-mCherry plasmid (modified gift from the Roy laboratory, Kaether et al., 2000) or 0.10 μg of clathrin-mCherry plasmid (gift from the Roy laboratory, Ganguly et al., 2021). Experiments were performed the next day, 16-24 h after transfection.

#### Pharmacological manipulations

Neurons were treated with either dimethylsulfoxide (DMSO) 0.1% (from pure DMSO, Sigma-Aldrich #D2650) or swinholide A 100 nM (from 100 µM stock in DMSO, Sigma-Aldrich #S9810) for 3 h in their original culture medium at 37°C, 5% CO2 and then transferred in Tyrode’s solution (119 mM NaCl, 25 mM HEPES, 2.5 mM KCl, 2 mM CaCl2, 2 mM MgCl2, 30 mM glucose, pH 7.4) complemented with either DMSO or swinholide A for live-cell imaging, or directly fixed and immunostained for SIM microscopy.

#### Fixation and immunostaining

Neurons were fixed and labeled as previously described (Jimenez et al., 2020). Briefly, neurons were fixed in freshly prepared 4% PFA in PEM buffer (80 mM PIPES pH 6.8, 5 mM EGTA, 2 mM MgCl2) for 10 min at 35°C. After rinsing 3 times in 0.1 M phosphate buffer (PB), neurons were blocked for 2-3 h at RT in immunocytochemistry buffer (ICC: 0.22% gelatin, 0.1% Triton X-100 in PB), and incubated with primary antibodies diluted in ICC for 2 h at room temperature or overnight at 4°C. After rinsing 3 times in ICC, neurons were incubated with secondary antibodies diluted in ICC for 1 h at RT and rinsed. Filamentous actin was labeled post-immunostaining using phalloidin Atto643 for 2 h at RT. Stained coverslips were kept in PB complemented with 0.02% sodium azide at 4°C until imaging by SMLM. For SIM, coverslips were mounted in ProLong Glass (Thermo Fisher Scientific #P36980).

#### Antibodies labeling

| Primary antibodies and labels | Dilution | Source |
| --- | --- | --- |
| Chicken polyclonal anti-Map2 | 1:1000 | Synaptic Systems (#188 006) |
| Sheep polyclonal anti-GFP | 1:500 | Bio-Rad (#4745 1051) |
| Guinea pig monoclonal anti-synap-<br>tophysin | 1:400 | Synaptic Systems (#101 308) |
| Rabbit polyclonal anti-βIV-spectrin<br>(against residues 2237–2256 of<br>human βIV-spectrin) | 1:500 | gift from Matthew<br>Rasband<br>(Houston, USA) |
| Mouse monoclonal anti-βII-spectrin<br>(clone 42 against residues 2101–<br>2189 of human β2-spectrin) | 1:100 | BD Biosciences<br>(#612563) |
| Mouse monoclonal anti-αII-spectrin<br>(clone D8B7) | 1:100 | Biolegend (#803201) |
| Mouse monoclonal anti-syntaxin 1A | 1:200 | Synaptic Systems<br>(#110011) |
| Rabbit polyclonal anti-clathrin heavy<br>chain | 1:150 | Abcam (Ab 21679) |
| Phalloidin Atto643 | 1:40 | AttoTech (AD 643-81) |

| Secondary antibodies | Dilution | Source |
| --- | --- | --- |
| Donkey anti-chicken DL405 | 1:100 | Jackson (703-475-155) |
| Donkey anti-sheep AF488 | 1:400 | Abcam (Ab 150177) |
| Donkey anti-mouse AF488 | 1:400 | Invitrogen (A21202) |
| Goat anti-guinea pig AF555 | 1:400 | Invitrogen (A21435) |
| Donkey anti-rabbit AF555 | 1:400 | Invitrogen (A31572) |
| Donkey anti-mouse AF647 | 1:300 | Invitrogen (A31571) |
| Donkey anti-rabbit CF680 | 1:150 | Biotium (BTM20820) |
| Rabbit polyclonal anti-clathrin<br>heavy chain | 1:150 | Abcam (Ab 21679) |

#### Live-cell AIS staining

Live-cell AIS staining was performed as previously described (Farías et al., 2016) using a mouse anti-pan-Neurofascin (external) antibody (clone A12/18, Antibodies Incorporated #75-172) targeting an extracellular epitope of neurofascin 155/186 conjugated with CF647 using a Mix-N-Stain CF647 antibody labeling kit (Sigma-Aldrich #MX647S50) following the manufacturer protocol. The conjugate was diluted to 1:50 and incubated for 10 min at 37°C and 5% CO2 in the original culture medium, and subsequently briefly washed in Tyrode’s solution before live-cell imaging.

### Microscopy

#### Live-cell microscopy

Live-cell imaging of neurons expressing vamp2-pHluorin and synaptophysin-mCherry or clathrin-mCherry was performed on an EclipseTi2-E inverted microscope (Nikon Instruments) equipped with an ORCA-Fusion sCMOS camera (Hamamatsu Photonics K.K. - C14440-20UP) and a CFI SR HP Apochromat TIRF 100X Oil (NA 1.49) objective. The system was equipped with a Nikon Perfect Focus System (PFS) and images were acquired with the NIS-Elements AR 5.30.05 software. Coverslips with neurons were mounted in a Ludin metal chamber (Life Imaging Services) in Tyrode’s solution. Neurons were maintained in a humid chamber at 35.5-37°C for the duration of the experiments using a stage-top incubator (Okolab inc). Illumination was performed in HiLo mode, i.e. by setting the laser incidence at the critical angle to generate a ~1 µm light sheet at the surface of the coverslip, resulting in rejection of out-of-focus fluorescence and better SNR. To image exocytosis, vamp2-pHluorin present initially at the plasma membrane was photobleached by exposing it to high power 488 nm laser light (100%) for 30 s before time-lapse imaging to remove the basal membrane signal and highlight exocytic signal (Yudowski et al., 2007). 200 ms exposure frames were then continuously acquired for 90 s using 488 nm laser light at lower power (1-10 %). To identify synapses, synaptophysin-mCherry was imaged before the detection of exocytosis using 561 nm laser light at 1-2% with 200 ms exposure.

For 2-color imaging of vamp2-pHluorin and clathrin-mCherry, the photobleaching time of vamp2-pHluorin was reduced to 15 seconds, the vamp2-pHluorin 488 nm illumination to 5% for 15 s before a single image of clathrin-mCherry was captured using the 561 nm laser light at 15-20% with 500 ms exposure. This sequence of 15 s of vamp2-pHluorin acquisition and a single frame of clathrin-mCherry illumination was repeated for 5 minutes.

#### Structured illumination microscopy (SIM)

SIM of neurons was performed on an N-SIM-S microscope (Nikon Instruments). The N-SIM system uses an inverted Nikon Eclipse Ti2-E microscope with a perfect focus system, an integrated Nikon laser launch with 405, 488, 561, and 640 nm solid-state excitation lasers, a 100X NA 1.49 oil objective (SR HP Apo TIRF), a Mad City Labs Nanodrive piezo stage and a Hamamatsu Fusion BT CMOS camera. On fixed cells, after locating a neuron of interest using low-intensity illumination, a z-stack was acquired in 2D-SIM (9 images per image) mode with a step-size of 0.12 μm. Reconstruction was performed using the N-SIM module in the NIS-Elements software (AR 5.30.05), resulting in a ~120 nm lateral resolution and images were reconstructed into a maximum projection.

#### Correlative live - cell and SMLM acquisition and post-processing

During the live-cell imaging session, an objective-style diamond tip (Leica) was used to scratch a line under the coverslip. A 40x objective was used to locate the ends of the scratched line and store their XY coordinates in the NIS software, before storing the positions of the imaged neurons and acquiring a reference phase image for each. After the live-cell imaging session, neurons were immediately fixed, and proteins of interest were labeled using immunocytochemistry. Meanwhile, the vamp2-pHluorin timelapse movies were analyzed to identify sites of exocytic events, to select the corresponding axonal segments for SMLM imaging. Stained coverslips were transferred to the SMLM setup (see below). The line ends of the coverslip scratch were located using a 40X objective and stored. Using a custom Fiji macro, the coordinates of the imaged neurons on the live-cell setup stage were transformed into coordinates on the SMLM setup stage using the stored positions of the scratch ends for each setup, generating a stage positions file that was loaded on the SMLM setup and used to relocate the imaged neurons. The axonal segment of interest with detected exocytic events was then manually relocated before acquisition of the SMLM image.

#### Image acquisition for spectral demixing STORM

Fixed, immunostained cells were secured to a silicone perfusion chamber (Electron Microscopy Sciences, USA) filled with reducing imaging buffer (Smart Kit Buffer A and glucose oxidase, Abbelight, France; 2-mercaptoethylamine (MEA) diluted in 360mM HCl, Sigma #30070) fixed to a glass slide (Jimenez et al., 2020). Prepared samples were mounted on a Nikon Eclipse Ti2 inverted microscope body (Nikon) equipped with a motorized stage, a piezo Z-stage (Mad City Labs), a focus stabilization system (Perfect Focus System, Nikon), and a high numerical aperture 100X oil-immersion objective lens (1.49 NA, CFI SR HP Apochromat TIRF 100XC, Nikon). HiLO illumination was used to restrict illumination to ~1 μm above the glass interface. Illumination was provided by two 640 nm continuous wave diode lasers (Oxxius, France) with a combined power between 300 and 400 mW measured at the back aperture. An ASTER module (Abbelight, France) scanned the beam in a 70 µm x 70 µm raster (Mau et al., 2021), yielding a flat-top square illumination profile at the sample plane with a power density between 6.12 and 8.16 kW cm-2 in the imaged region of interest. Fluorophores in long-lived dark states were recovered with a 405 nm continuous wave diode laser in the same illumination beam line, at powers <15 mW measured at the back aperture. The microscopy body, stage positions and filter wheel were controlled using NIS-Elements software (version 5.42.02, Nikon), while cameras and illumination lasers for SMLM acquisitions were controlled using NEO Live imaging software (version 2.16.3, Abbelight). Diffraction-limited epifluorescence images were acquired using an inline LED light source (CoolLED).

Fluorescence emission was detected in a split light path optimized for spectral demixing (Friedl et al., 2023). Briefly, emission signal was isolated with a quad-edge dichroic beam-splitter (Di03-R405/488/532/635-t3-25×36, Semrock) and stray light filtered further on a quad-band bandpass filter (FF01-446/510/581/703-25, Semrock). Astigmatic shaping of the PSF was achieved with a cylindrical lens, and a dichroic mirror (FF699-Fdi01-t3-25×36) divided the filtered emission into reflected and transmitted image paths, detected by two water-cooled 16 bit sCMOS cameras (ORCA-Fusion BT, Hamamatsu Photonics). Acquisition sequences of 512 x 512 pixels (pixel size 97 nm) were recorded for 60,000 frames at an exposure of 20 ms.

### Data quantification and analysis

#### Timelapse movies preprocessing

Acquired videos were preprocessed using the Filter Timelapse script (available at https://sites.imagej.net/Christophel-eterrier/plugins/NeuroCyto%20Lab/Kymographs/) that performed image stabilization using the Image Stabilizer plugin (https://imagej.net/plugins/image-stabilizer) after 2X downscaling, and bleach correction using average intensity compensation of the foreground identified as the top 12% of pixel intensities within each image of the sequence.

#### Exocytosis detection

Preprocessed videos of vamp2-pHluorin transfected neurons were analyzed using the pHusion script written in ImageTank (O’Shaughnessy et al., 2024). Briefly, exocytotic events were detected by using the Difference of Gaussian methods using a sigma of 3 and a scaling factor of 1.25. Using the region of interest from the DoG image, a small image series was cropped from the raw image stack taking 5 frames before and 30 after the identified spot. Each image in the series was fitted with a gaussian model. Only events with a R2 (goodness-of-fit) for the gaussian model greater than 0.7 were considered. Events that drifted greater than 4 pixels were removed. Identified exocytosis events were subsequently manually verified in Fiji. Identified exocytosis events were manually classified as occurring either within synapses or on axonal shaft segments using the synaptophysin-mCherry as a visual guide.

#### Data processing for spectral demixing STORM

Paired single molecule localizations were detected in the reflected and transmitted images using the globLoc fitting algorithm (Li et al., 2022), incorporated in the ‘fit_global_dual-channel’ module in the Super-resolution Microscopy Analysis Platform (SMAP)(Ries, 2020). Candidate peaks were initially detected in each channel separately, with a dynamic factor of 1.7 and a minimum distance of 7 pixels between peaks. The global localization step was performed on a CUDA enabled NVIDIA GeForce RTX 3090 graphics processor, with a dual-channel cubic spline interpolated PSF model iterated up to 150 times on the candidate peaks to optimize Cramer-Rao bound localization statistics. Localization coordinates were subsequently manually classified into 2 channels according to the ratiometric distribution of photon intensity detected in the reflected and transmitted images. Drift correction was calculated using FFT-based cross-correlation on block-wise time series of the raw localizations. Positional aberrations introduced by the cylindrical lens astigmatism were corrected for by projective transformation of the localization coordinates, using a transform matrix which was previously calibrated from images of fluorescent beads both with and without the cylindrical lens in place. Localization coordinates were reconstructed as images at 16 nm using the ImageJ ThunderSTORM plugin (Ovesný et al., 2014) with thresholding below 5 detections to remove spurious signals with long-lived on states, and thresholding below 30 nm uncertainty in lateral precision.

#### Correlative live-cell and SMLM analysis

Initially, an RGB merge of the HiLo vamp2-pHluorin image and the phase construct image from the live-cell data (allowing the appearance of non-transfected cellular structures) was aligned to an RGB merge of the corresponding high-magnification SMLM image of αII/βII-spectrin (labeling the majority of axonal structures in the field of view) using the Fiji plugins BigDataViewer (Pietzsch et al., 2015). The resulting affine transform was then applied to the live-cell vamp2-pHluorin data using TransformJ (Meijering et al., 2001) to align them to the SMLM reconstructions (at 16 nm per pixel). The vamp2-pHluorin movies were up-sampled using a quintic B-spline approach. Affine transformed Hilo images were then manually cropped to the size of the SMLM image.

#### Structural repetition analysis of axon segments

The approach used to map periodic ring structures closely followed the method described in (Mendes et al., 2025) using ImageJ/Fiji functions, plugins and custom macros (available at https://www.github.com/HenriquesLab/SReD). First, we estimated the local orientations of axons by applying a Gaussian blur (20 px sigma) to the images, which preserved high-order structures while suppressing single-molecule, low-order information. We then applied Otsu’s thresholding (Otsu, 1979) to remove unwanted objects from the images and produce binary masks. Next, we used the “Skeletonize” function on the thresholded images to extract 1 pixel-wide axon skeletons. To minimize diagonal discontinuities inherent to these thin lines, we applied an additional Gaussian blur (2 px sigma) to the skeletons. All skeleton images were then range-normalized to a common intensity interval. To analyze orientation-specific repetition, we generated a set of 45×45 pixel image blocks, each containing a single 1 pixel-wide line oriented from 0º to 180º in 10º increments. These blocks were also Gaussian blurred (sigma = 2 px) to match the appearance of the axon skeletons. We then applied SReD’s block repetition method, using the Pearson’s correlation coefficient and a relevance constant of 0, to compare the line blocks with the axon skeletons and generate repetition maps. In these maps, regions containing skeletons aligned with the tested orientations exhibited higher Structural Repetition Score (SRS). Each repetition map was normalized, multiplied by the corresponding Gaussian-blurred mask to exclude unwanted detection, and normalized again to standardize intensity.

To detect ring patterns, we used a reference block containing three vertical lines, mimicking a side-view 2D projection of rings. This 3-ring pattern was defined by two parameters: inter-ring spacing (192 nm, previously optimized using similar data sets (Mendes et al., 2025; Vassilopoulos et al., 2019) and ring height (chosen to span approximately one third of the axon girth, enabling detection of local rather than global patterns). For each angle (0º to 180º in 10º increments), input axon images were duplicated, zero-padded to prevent border cropping, and rotated accordingly. Each rotated image was range-normalized before analysis. Block repetition maps were then calculated using the optimized ring block, the Pearson correlation coefficient, and a relevance constant of 0. After computation, repetition maps were rotated back to their original orientation. The repetition maps were then rotated and cropped to match the original image dimensions. Previously computed orientation maps were Gaussian blurred (20 px sigma) to encompass full axon segments and used as orientation masks for their corresponding repetition maps, ensuring each map retained only angle-specific information. To further enhance specificity, a power transformation (exponent 2) was applied to the final repetition maps. A complete view of all the ring patterns detected was generated using the “Z Project > Sum of slices” function.

#### Line profiles of clathrin pits, exocytosis events, spectrin and repetition maps

To identify CCPs and remove clathrin signals not associated with CCPs, super-resolved images were intensity-thresholded and used as masks. The repetition maps provided by SreD were normalized between 0 and 1 and for each given line profile the envelope was calculated each 160 nm. Line profiles were manually aligned to the center of the exocytosis site. Obtained line profiles were finally normalized between 0 and 1.

#### Statistics

Individual measurement points (neurons) from independent experiments (cultures) were pooled. Profiles, graphs, and statistical analyses were generated using Prism (GraphPad). Normality and outliers were determined using the Shapiro-Wilk test and Rout methods, respectively. Significances were tested either using Mann-Whitney test (Figure 1F, 3B-D) or oneway, non-parametric ANOVA with Kruskal-Wallis post-hoc significance testing (Figure 2C). In Figures, the results of the post-hoc significance are indicated as follows: ns or ns, non-significant; ^*^, p<0.05; ^**^, p<0.01; ^***^, p<0.001. On bar graphs, dots are individual data points of each independent experiment, bars or horizontal lines represent the mean, and vertical lines are the SEM unless otherwise specified. Boxplots show the first to third quartile with whiskers ranging from 5-95% of the data distribution. Statistics are summarized in the Supplementary Data file.

## Supplementary Material

**Figure S1:**
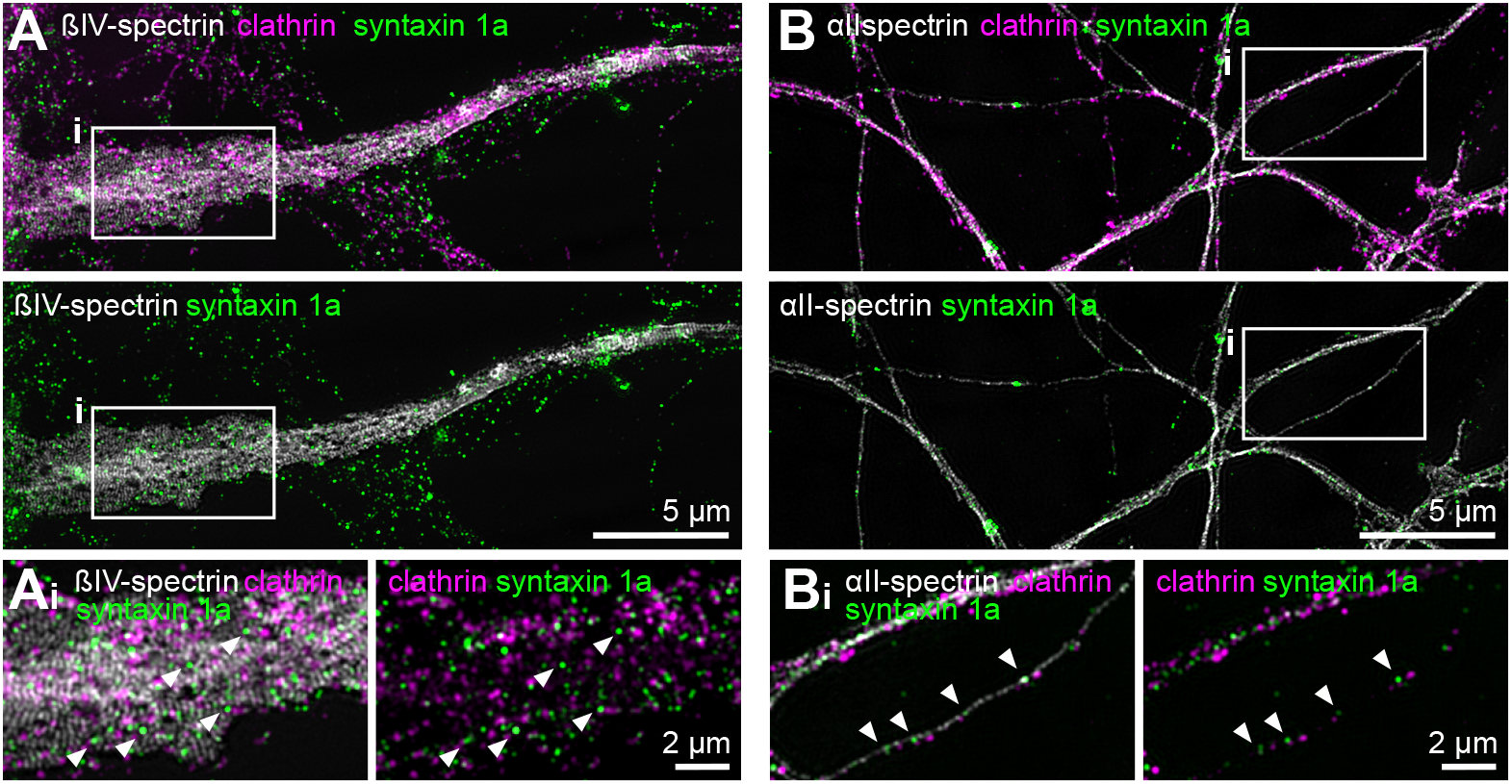
Distribution of syntaxin 1a along axons. Representative images acquired using structured illumination microscopy (SIM) of syntaxin 1a (green) in either the AIS (A) with βIV-spectrin (grey) and clathrin (magenta, top only), or in the distal axon (B) with αII-spectrin (grey) and clathrin (magenta, top only). White arrowheads show syntaxin 1a clusters at the axonal shaft.

**Figure S2:**
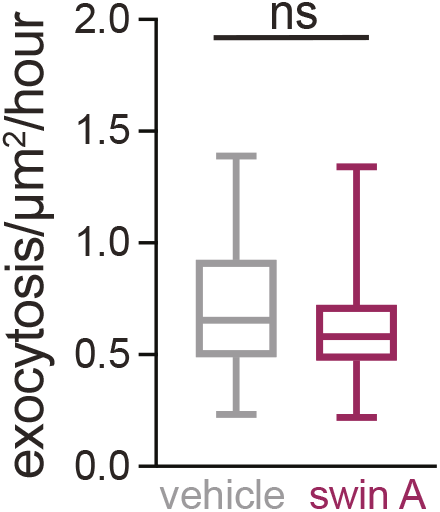
Exocytosis rate along the whole distal axon (both synapse and axonal shaft). Exocytosis rate along the distal axon (both synapse and axonal shaft). Fields of view = 29, 33; 5 independent cultures, Mann-Whitney test.

**Figure S3:**
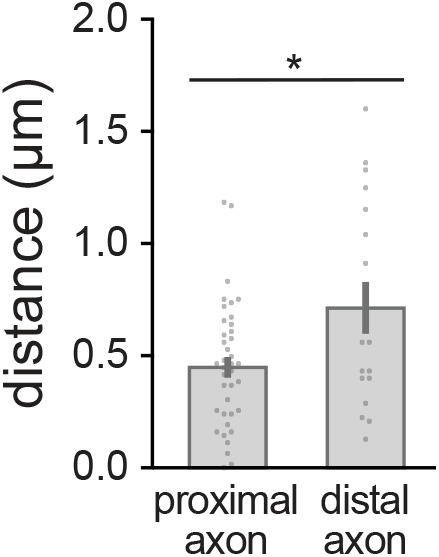
Spatial separation of exocytosis sites and surrounding CCPs in proximal and distal axons. Distance between exocytic sites and nearest CCPs in proximal (exocytic events = 39, 4 neurons, 4 independent cultures) and distal axon (exocytosis events = 17, 4 neurons, 2 independent cultures, unpaired t-test).

**Movie S1:**
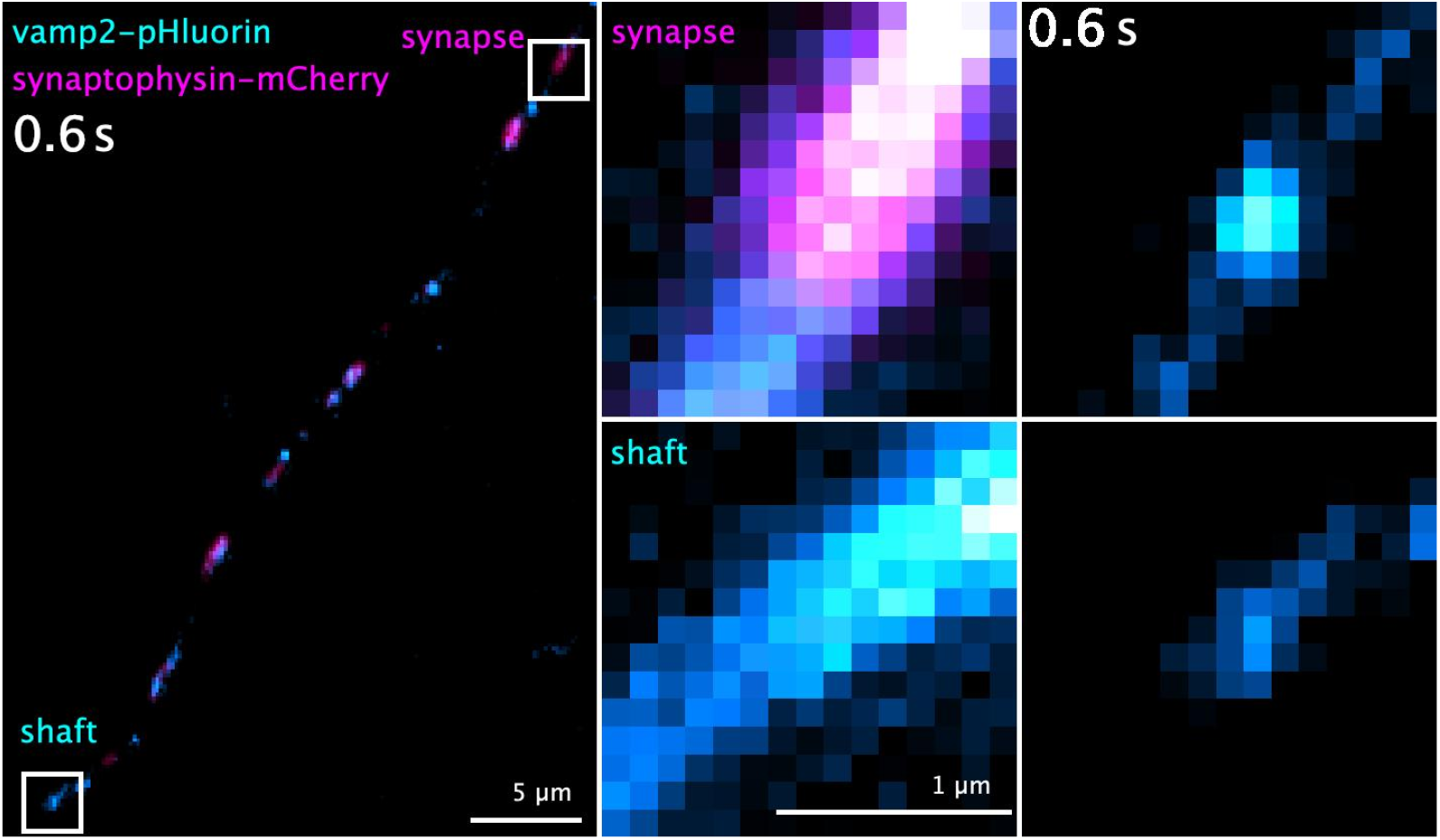
Exocytosis occurring at a presynapse and along the axonal shaft. Left, timelapse (one frame every 0.2 s for 9.4 s) of an axon from a hippocampal neuron expressing vamp2-pHluorin and synaptophysin-mCherry, imaged using HiLo microscopy. Right, timelapse (one frame every 0.2 s for 2 s) corresponding to exocytosis occurring at synapse (top) and axonal shaft (bottom) corresponding to the white rectangles shown on the left.

**Movie S2:**
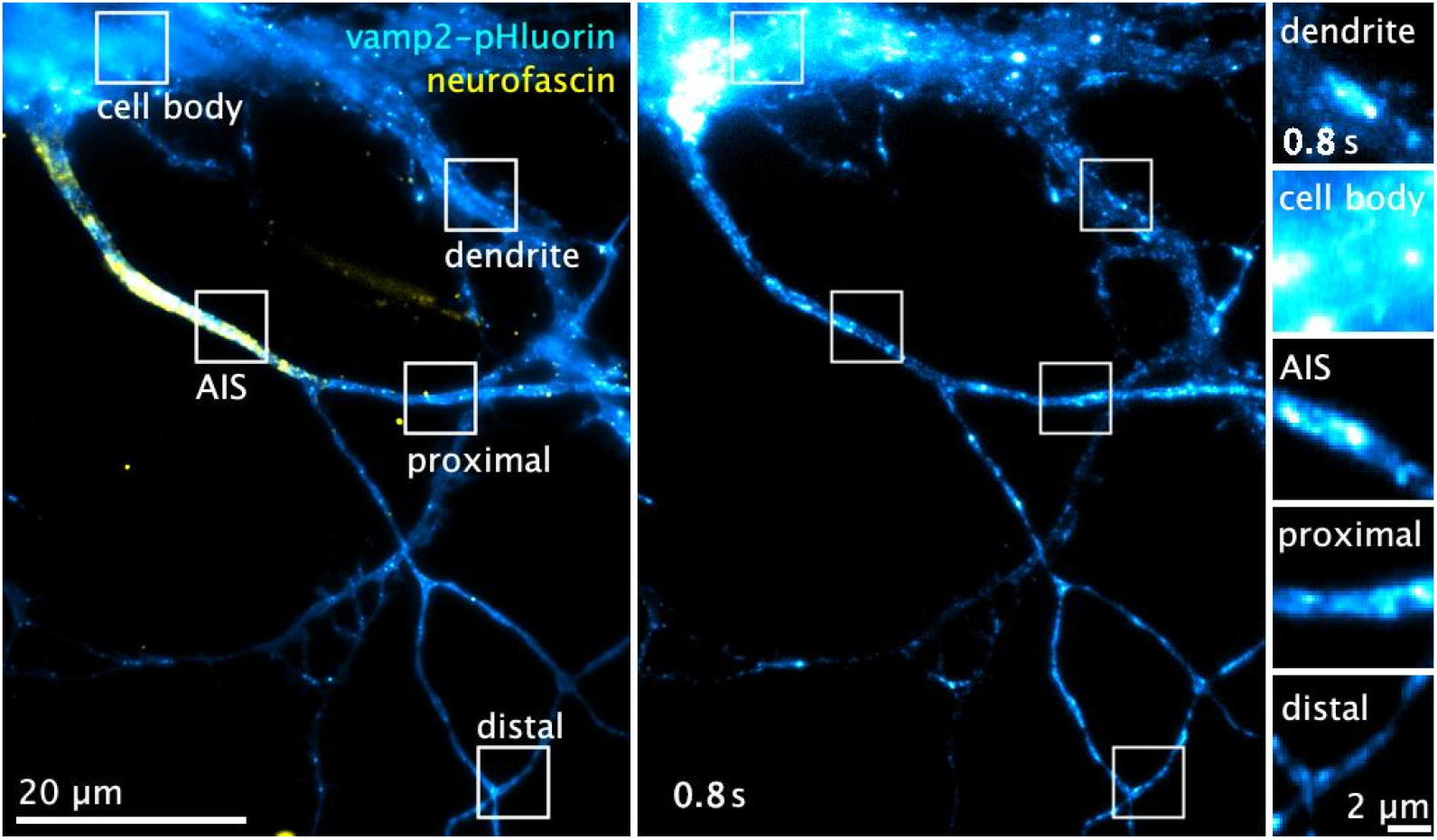
Examples of exocytic events in different neuronal compartments. Left, hippocampal neuron in culture expressing vamp2-pHluorin (cyan-hot) with the AIS labeled for neurofascin (yellow), imaged using HiLo microscopy. White rectangles highlight exocytic sites occurring in distinct compartments: cell body, dendrites, AIS, proximal and distal axon. Middle, timelapse of vamp2-pHluorin for this neuron (one frame every 0.2 s for 90 s). Right, timelapse of the regions of interest corresponding to the white rectangles shown on the left (one frame every 0.2 s for 2 s).

**Movie S3:**
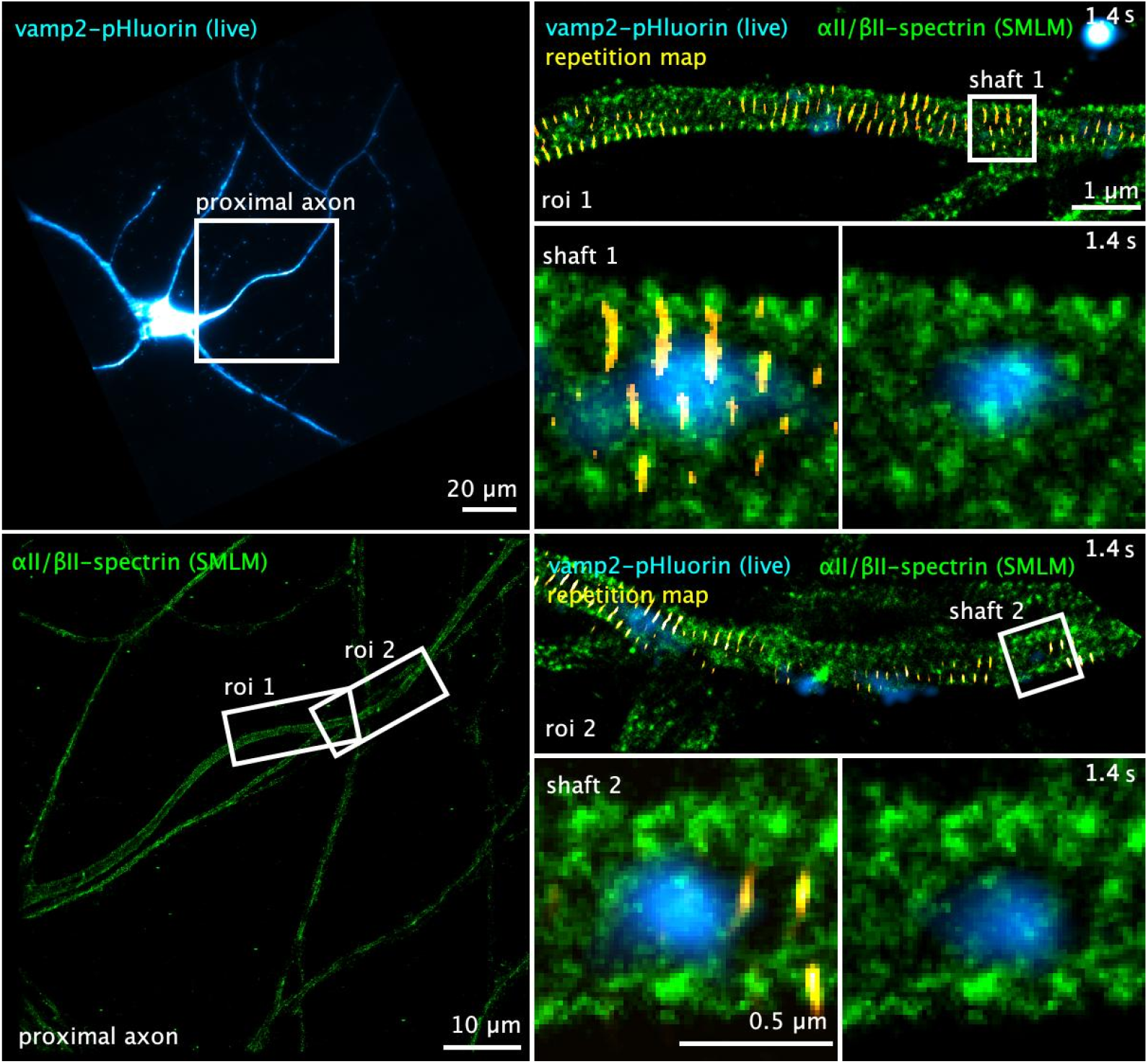
Nanoscale organization of αII/βII-spectrin at exocytic sites along the proximal axon. Top left, hippocampal neuron in culture expressing vamp2-pHluorin (cyan-hot) imaged using HiLo microscopy. Image is affine transformed. White rectangle highlights the field of view of the SMLM acquisition. Bottom left, αII/βII-spectrin (green) nanoscale organization obtained by SMLM. White rectangles show the regions of interest shown on the right. Right, timelapse of the regions of interest 1 (top) and 2 (bottom) (one frame every 0.2 s for 10 s) of vamp2-pHluorin (cyan-hot) aligned to the αII/βII-spectrin (green) signal and the repetition map (yellow). White rectangle highlights the exocytotic site shown below, with a timelapse of the exocytic site at shaft 1 and 2 (one frame every 0.2 s for 3 s).

**Movie S4:**
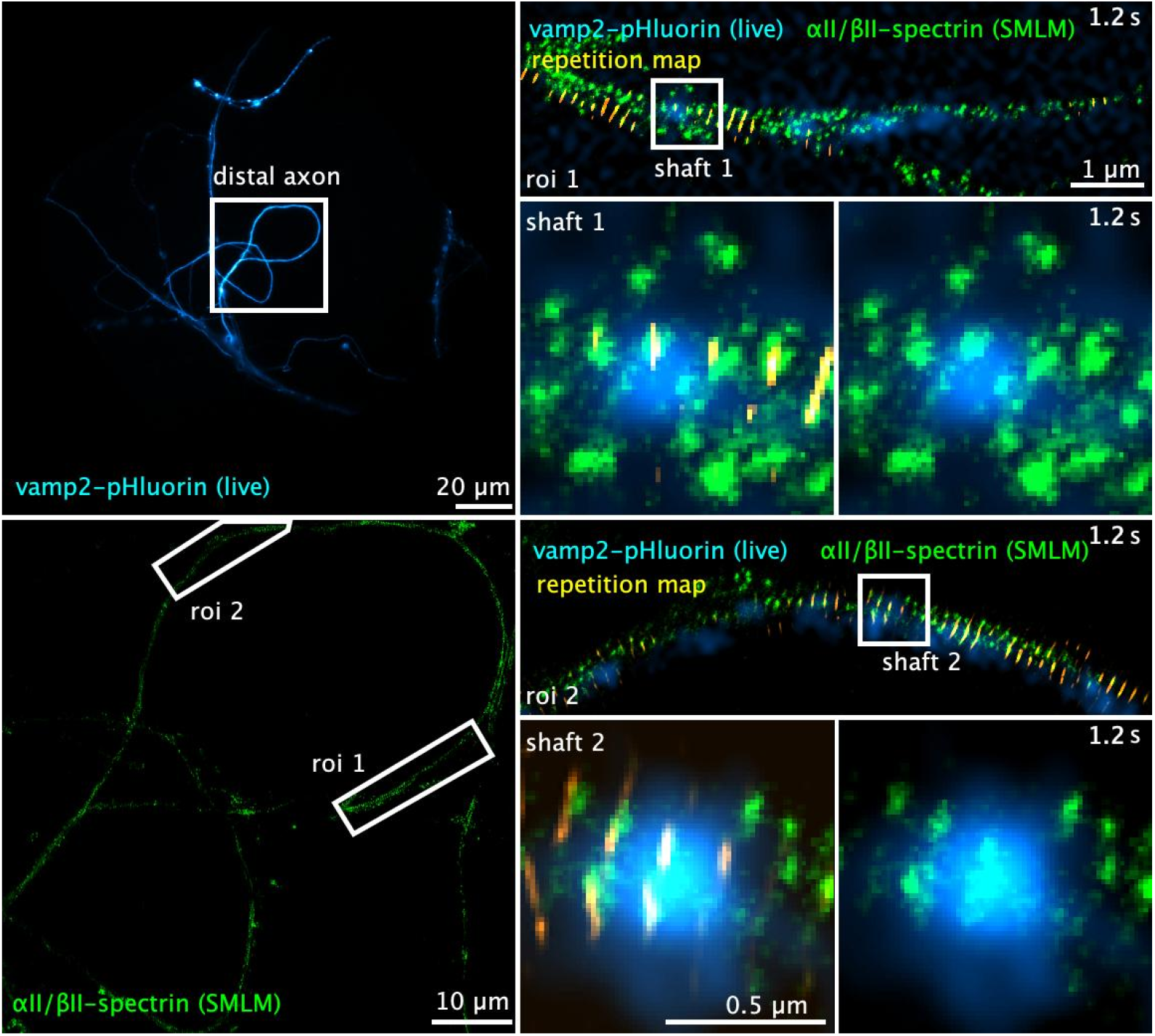
Nanoscale organization of αII/βII-spectrin at exocytic sites along the distal axon. Top left, hippocampal neuron in culture expressing vamp2-pHluorin (cyan-hot) imaged using HiLo microscopy. Image is affine transformed. White rectangle highlights the field of view of the SMLM acquisition. Bottom left, αII/βII-spectrin (green) nanoscale organization obtained by SMLM. White rectangles show the regions of interest shown on the right. Right, timelapse of the regions of interest 1 (top) and 2 (bottom) (one frame every 0.2 s for 10 s) of vamp2-pHluorin (cyan-hot) aligned to the αII/βII-spectrin (green) signal and the repetition map (yellow). White rectangle highlights the exocytotic site shown below, with a timelapse of the exocytic site at shafts 1 and 2 (one frame every 0.2 s for 3 s).

**Movie S5:**
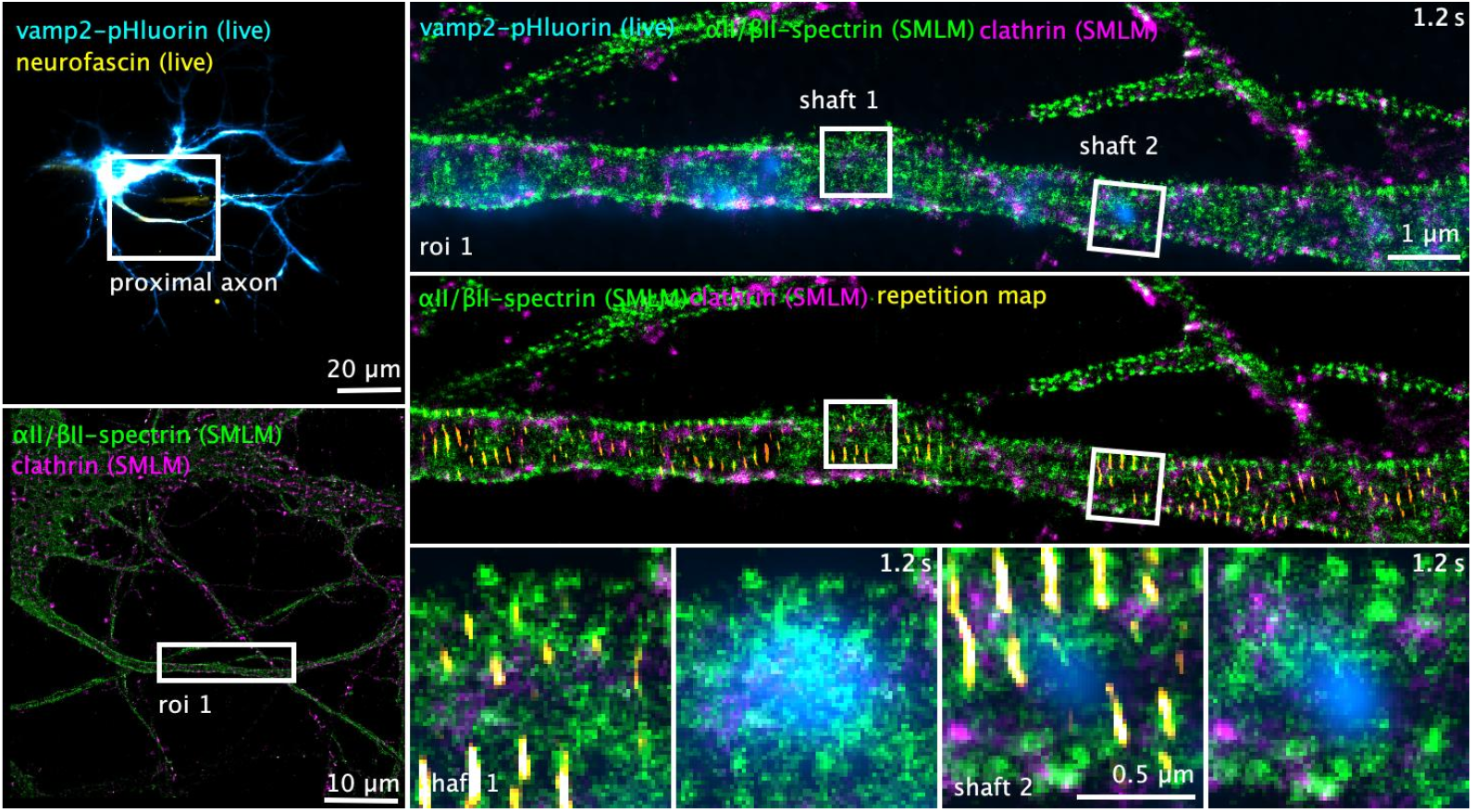
Nanoscale organization of αII/βII-spectrin and clathrin at exocytic sites along the proximal axon. Top left, hippocampal neuron in culture expressing vamp2-pHluorin (cyan-hot) imaged using HiLo microscopy. Image is affine transformed. White rectangle highlights the field of view of the SMLM acquisition. Bottom left, αII/βII-spectrin (green) and clathrin (magenta) nanoscale organization imaged by 2-color demixing STORM. White rectangle shows the regions of interest shown on the right. Top right, timelapse of the region of interest 1 (one frame every 0.2 s for 10 s) of vamp2-pHluorin (cyan-hot) aligned to the αII/βII-spectrin (green) and clathrin (magenta) signal and (right-middle) the repetition map (yellow). White rectangles highlight the exocytotic sites shown below, with a timelapse at the exocytic site (one frame every 0.2 s for 3 s).

**Movie S6:**
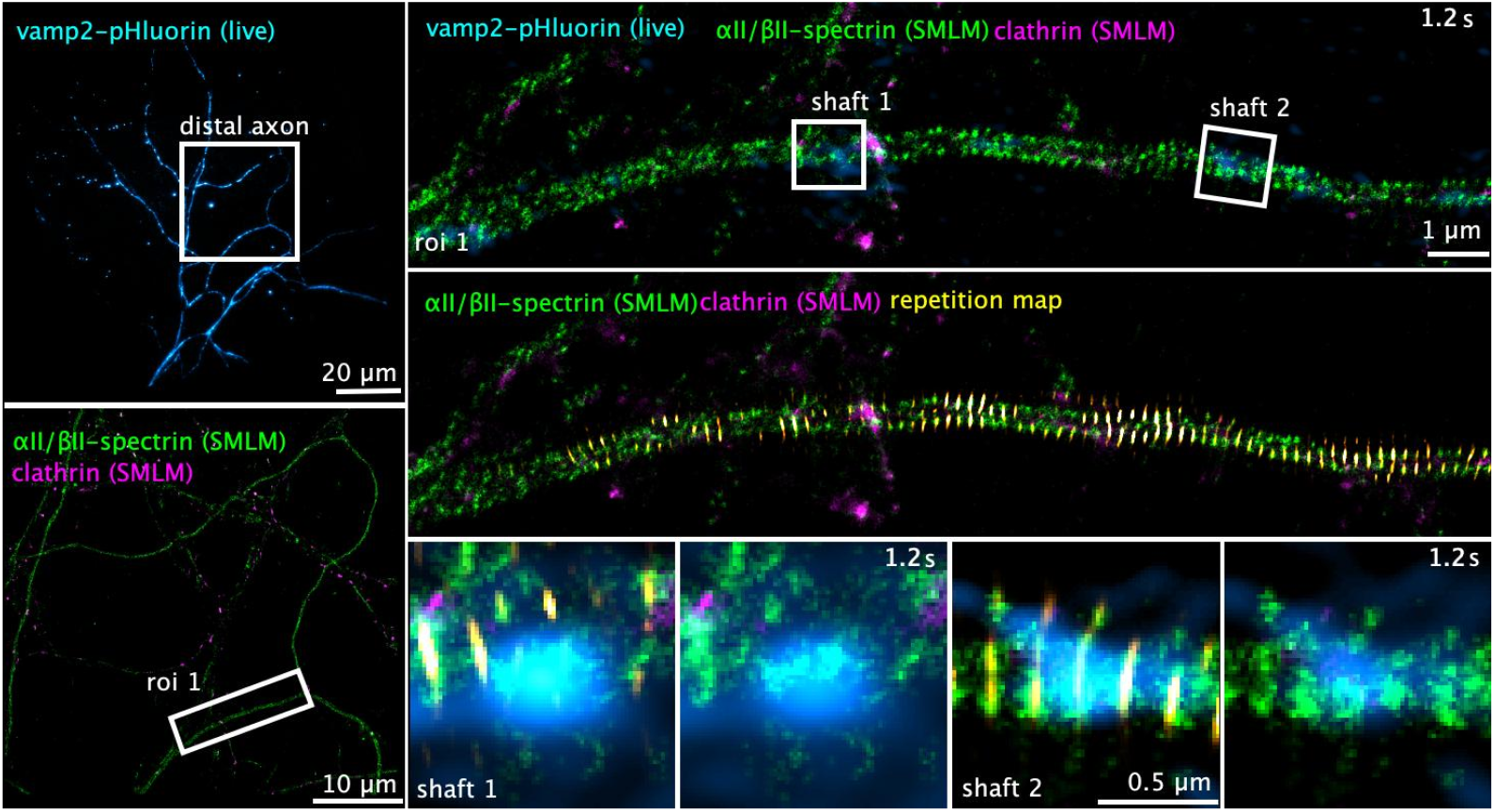
Nanoscale organization of αII/βII-spectrin and clathrin at exocytic sites along the distal axon. Top left, hippocampal neuron in culture expressing vamp2-pHluorin (cyan-hot) imaged using HiLo microscopy. Image is affine transformed. White rectangle highlights the field of view of the SMLM acquisition. Bottom left, αII/βII-spectrin (green) and clathrin (magenta) nanoscale organization obtained by 2-color demixing STORM. White rectangle shows the regions of interest shown on the right. (Right-top) Timelapse of the region of interest 1 (one frame every 0.2 s for 10 s) of vamp2-pHluorin (cyan-hot) aligned to the αII/βII-spectrin (green) and clathrin (magenta) signal and (right-middle) the repetition map (yellow). White rectangles highlight the exocytotic sites shown below, with a timelapse at the exocytic site (one frame every 0.2 s for 3 s).

**Movie S7:**
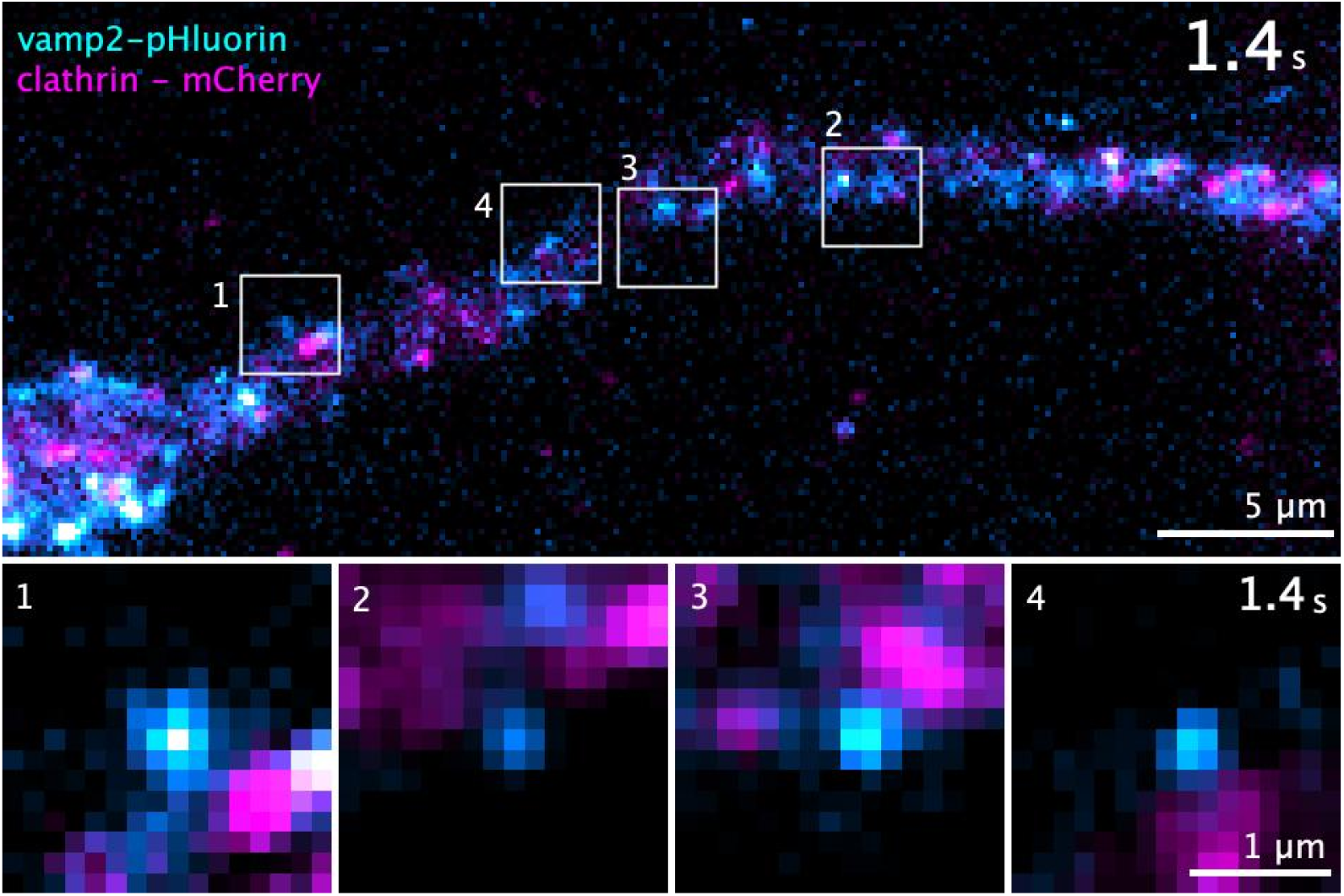
Sequential live-cell imaging of exo- and endocytosis along the AIS. Top, timelapse of the AIS of a hippocampal neuron in culture expressing vamp2-pHluorin (cyan-hot) and clathrin-mCherry (magenta) (one frame of vamp2-pHluorin every 0.2 s for 160 s interleaved with one frame of clathrin mCherry every 15 s). White rectangles show the timelapse regions of interest below. Bottom, timelapse (one frame of vamp2-pHluorin every 0.2 s for 3 s interleaved with one frame of clathrin mCherry every 15 s) of the regions of interest above.

## Supplementary Data

See the provided ***SupplementaryData*.*xlsx*** Excel file listing values, statistics and singificance tests used for all graphs shown in this manuscript.

## Notes

### Competing Interest Statement

The authors have declared no competing interest.

